# Embryonic lethality in mice lacking Trim59 due to impaired gastrulation development

**DOI:** 10.1101/171603

**Authors:** Xiaomin Su, Chenglei Wu, Xiaoying Ye, Ming Zeng, Zhujun Zhang, Yongzhe Che, Yuan Zhang, Lin Liu, Yushuang Lin, Rongcun Yang

**Affiliations:** Department of Immunology, Nankai University School of Medicine, Nankai University, Tianjin, China; Key Laboratory of Bioactive Materials Ministry of Education, Nankai University, Tianjin, China; State Key Laboratory of Medicinal Chemical Biology, Nankai University, Tianjin, China; College of Life Sciences, Nankai University, Tianjin, China; Shandong Provincial Key Laboratory of Animal Cells and Developmental Biology, Shandong University, Shandong, China

**Author notes:** These authors contributed equally to this work (Xiaomin Su, Chenglei Wu, Xiaoying Ye). Correspondence and requests for materials should be addressed to Rongcun Yang or Yushuang Lin or).

**Keywords:** Trim59, Embryonic lethality, Gastrulation, F-actin assembly, Ubiquitination

## Abstract

TRIM family members have been implicated in a variety of biological processes such as differentiation and development. We here found that Trim59 plays a critical role in early embryo development from blastocyst stage to gastrula. There existed delayed development and empty yolk sacs from embryonic day (E) 8.5 in *Trim59* -/- embryos. No viable *Trim59* -/- embryos were observed beyond E9.5. Trim59 deficiency affected primary germ layer formation at the beginning of gastrulation. In *Trim59* -/- embryos at E6.5 and E7.5, the expression of primary germ layer formation associated genes including *Brachyury*, *lefty2*, *Cer1*, *Otx2*, *Wnt3* and *BMP4* was reduced. Homozygous mutant embryonic epiblast was contracted and the mesoderm was absent. Trim59 could interact with actin and myosin associated proteins. Trim59 deficiency disturbed F-actin polymerization during inner cell mass differentiation. Trim59 mediated polymerization of F-actin was via WASH K63-linked ubiquitination. Thus, Trim59 may be a critical regulator for early embryo development from blastocyst stage to gastrula through modulating F-actin assembly.

## INTRODUCTION

Mouse embryological development is a dynamic process. It includes a series of cleavage divisions during development from embryonic day (E) 0.5 to E8.0 (Kojima et al., 2014). Following fertilization, one cell embryo (E0.5) undertakes a succession of cleavage divisions to generate a blastocyst (E4.5) which contains inner cell mass (ICM) and trophectoderm (TE) (Chazaud and Rossant, 2006). The ICM contains epiblast (EPI) precursor and primitive endoderm (PrE). From E5.5 to E6.0, pre-gastrula is formed during this stage. The formation of anterior visceral endoderm (AVE) represents a crucial event in the patterning of anterior-posterior (A-P) axis. Formation of the primitive steak (PS) marks the beginning of gastrulation at E6.5. During this stage, primary germ layers including ectoderm (EC), mesoderm (ME) and endoderm (EN) are formed (Tam et al., 2000).

During gastrulation, PS formation requires interactions between EPI, extra-embryonic AVE and extra-embryonic ectoderm (ExE) (Joubin and Stern, 1999). Multiple genes and signaling pathways are involved in the regulation of embryogenesis. In mouse embryos, EPI expresses *Nodal*; Whereas ExE expresses bone morphogenetic protein 4 (*BMP4*). These molecules are involved in many steps in pregastrulation development. With the growth of EPI, many of AVE genes are expressed in distal visceral endoderm (DVE) cells at E5.5. *Otx2* (Orthodenticle homeobox 2), *Cer1* (Cerberus 1) and *Lefty2* belong to AVE genes (Goncalves et al., 2011a). *Otx2* is a transcription factor; Whereas *Cer1* and *Lefty2* belong to signaling antagonists (Hoshino et al., 2015). Signaling pathways such as *β*-*catenin/Wnt3*, *Nodal*, *BMP4* and *FGF* are considered as essential to AVE formation (Soares et al., 2008). *Nodal* and *Wnt3* are expressed in the proximal epiblast adjacent to the extraembryonic ectoderm, next to where the PS arises (Liu et al., 1999); whereas *BMP4* is expressed in distal ExE (Padgett et al., 1993; Ray et al., 1991). At the same time, other posterior genes such as *Brachyury* and *Cripto* also play a critical role in the formation of primitive streak (PS) and definitive endoderm (DE) (Bassham and Postlethwait, 2000; Liguori et al., 2008; Takenaga et al., 2007).

Mouse embryological development is a complex developmental program with correcting mechanisms to avoid the transmission of errors (Arias and Hayward, 2006). However, the mechanism(s) of maintaining normal mouse embryological development is not completely understood. Multiple processes control cell lineage specification from blastocyst to three germ layer formation. These processes include cell-cell contact, gene expression, cell signaling pathways, positional relationships and epigenetics. However, actin remodeling may also be important for normal embryo development (Tan et al., 2015b). Studies have shown that cell actin skeleton is regulated by many proteins that either promote or inhibit actin polymerization during embryo development (Tan et al., 2015a).

In this study, we investigate the impact of Trim59 on the early embryonic development. Trim59, a member of the tripartite motif (TRIM)-containing protein superfamily, is characterized by one or two zinc binding motifs, an associated coiled-coil region and a RING-finger domain (Meroni and Diez-Roux, 2005). Previous studies show that Trim59 participates in many pathological regulation such as inflammation (Chen et al., 2017), cytotoxicity (Zhao et al., 2012), and especially tumorigenesis (Aierken et al., 2017; Khatamianfar et al., 2012; Luo et al., 2017; Zhang and Yang, 2017; Zhou et al., 2014). Here we find that Trim59 plays a vital role in mouse early embryonic development stage. Trim59 deficiency affects the formation of primary germ layers ectoderm, mesoderm and endoderm. Trim59 may promote F-actin assembly through WASH K63-linked ubiquitination during blastocyst develops into gastrula stage.

### Results

#### Trim59 deficiency causes early embryonic lethality

Trim59 could be detected not only in murine F1 ESCs (embryonic stem cells derived from C57BL/6J×C3H F1 mouse) but also in wild type (wt) murine E6.5-E9.5 embryos (Fig. S1), implying that Trim59 may play a role in embryonic development. To test this, we generated *Trim59* knockout (-/-) mice (Fig. S2). We observed more than six generations within a year and found that around 67% of the offsprings (70 animals) were *Trim59* +/- and 33% (34 animals) were *Trim59* +/+; whereas no offspring was *Trim59* -/- (Table 1), indicating that *Trim59* -/- genotype is embryonic lethality. To determine the embryonic stage of lethality, *Trim59* +/- mice were crossed and then embryos were dissected for genotyping. At E6.5, among 41 decidua examined, 12 (29%) were *Trim59* +/+, 20 (49%) *Trim59* +/- and 9 (22%) *Trim59* -/-, fitting well to a single-gene inheritance model, suggesting that *Trim59* knockout does not affect the implantation and decidualization but the development degree of *Trim59* -/- embryos was delayed at this stage (Table 1). Similar phenomenon was also observed at E7.5 (Table 1). At E8.5, some empty swollen decidua embryos could be observed in the dissected embryos. These abnormal embryos were *Trim59* -/- genotype (Table 1). Of the 57 offspring examined, 20 (35%) were empty, 11 (19%) contained embryos that were *Trim59* +/+ and 26 (46%) contained embryos that were *Trim59* +/- (Table 1). No *Trim59*-/- embryos were detected at E9.5 (Table 1). Taken together, these results indicate that Trim59 is essential for early embryonic development.

**Table 1.** Genotype analyses of offsprings fromTrini59+/- intercross.

| Stage | +/+ | +/- | -/- | Total | Phenotype |
| --- | --- | --- | --- | --- | --- |
| E6.5 | 12 | 20 | 9 | 41 | Delayed |
| E7.5 | 10 | 29 | 11 | 50 | Delayed |
| E8.5 | 13 | 24 | 12 | 49 | Empty yolk sacs |
| E9.5 | 11 | 26 | 0 (20*) | 57 | * Resorbent |
| Postnatal | 34 | 70 | 0 | 104 |  |

<sup>#</sup>Numbers indicate born mice or embryos detected at different stages of gestation

#### Trim59 deficiency affects gastrulation during early embryonic development

Due to some empty swollen decidua embryos were detected in the dissected embryos at E9.5 (Fig. 1A), Trim59 may affect the gastrulation which initiates from E6.5 and end at E7.5. H&E staining revealed that *Trim59* +/+ embryos exhibited normal morphology with the remodeling of epiblast, the formation of epithelium around a central lumen, which leads to the formation of postimplantation epiblast, and the emergence of primitive streak and mesoderm at E6.5 (Fig. 1B). At E7.5, embryo develops into late stage of gastrulation, aminion, aminion cavity, ExE cavity and intact three germ layers were observed in wt embryos; Whereas similar structure was not observed in *Trim59* -/- embryos. Conversely, contracted embryonic epiblast, absent mesoderm and collapsed skeleton often appeared in *Trim59* -/- embryos at E6.5 (Fig. 1B). Yolk sac and amnion were also defective in *Trim59* -/- embryos at E7.5 and no normal three germ layers was observed (Fig. 1B). These results suggest that Trim59 knockout affects gastrulation during early embryos development.

**Fig. 1.**
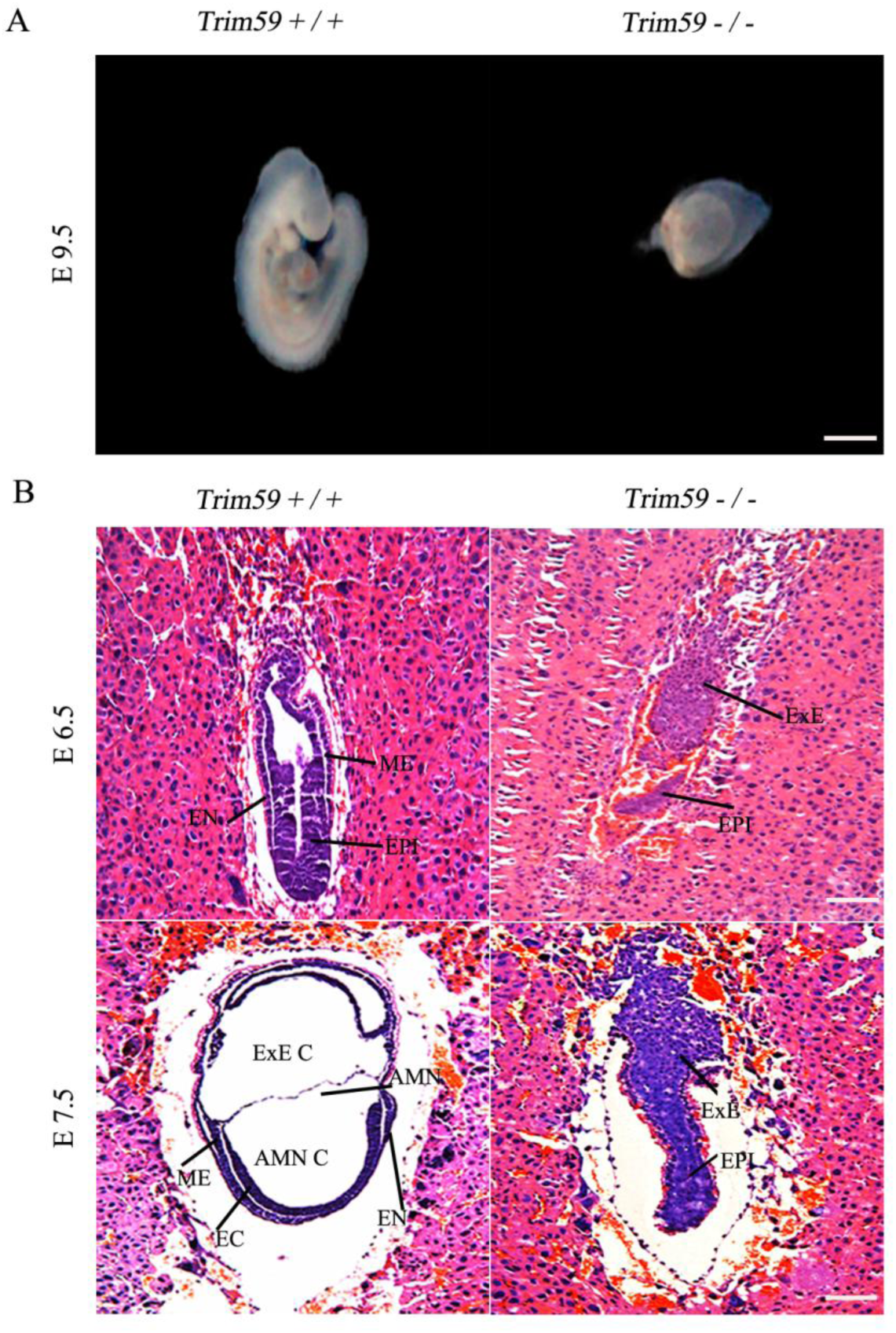
Trim59 is essential for normal embryogenesis. (A) Wild-type (*Trim59*+/+) and Trim59 knockout mice (*Trim59*-/-) embryos. Representative images of foetuses at E9.5 were displayed. (B) Hematoxylin and eosin staining of *Trim59*+/+ and *Trim59* -/- embryos at E6.5 and E7.5. Mesoderm (ME), entoderm (EN), ectoderm(EC), epiblast (EPI), extra-embryonic ectoderm (ExE), extra-embryonic coelom (ExE C), amnion (AMN), amnion cavity (AMN C). Scale bar, 100μm.

Next, we further characterized lethality stage of the early embryonic development. Formation of PS happens around E6.5, which originates from an elongated thickening of epiblast and marks the beginning of gastrulation (Gaivao et al., 2014). Many molecular markers are involved in the formation of PS including *Brachyury*, *Lefty2*, *Cer1* and *Otx2* such as that *Brachyury* marks the formation of PS and axial mesoderm (Caronna et al., 2013). We found that *Brachyury* could not be detected in *Trim59* -/- embryos (Fig. 2A-B). To exclude the possibility of a simple delay in the expression of *Brachyury*, we also tested its expression at E6.5 and E7.5. At neither stages did *Trim59* -/- embryos express *Brachyury* (Fig. 2A-B and E-F), indicating that Trim59 deficiency at early embryonic development stage fails to form PS. *Lefty2*, as a nascent mesoderm marker (Kim et al., 2014; Lenhart et al., 2011) could be detected at E6.5 in *Trim59* +/+ embryos, whereas *Trim59* -/- embryos were negative (Fig. 2C-D). These suggest that Trim59 is necessary for PS formation, which happens at early gastrulation. During gastrulation, Otx2 in the anterior neuroectoderm (ANE) (Simeone and Acampora, 2001) and Cer1 in the definitive endoderm (DEE) (Goncalves et al., 2011b) were markedly decreased in Trim59 knockout mice than in wt mice at E7.5 (Fig. 2G-H and I-J), representing failed development of ANE and DEE when Trim59 losses its function during gastrulation. These data support our findings that Trim59 affects gastrulation during early embryonic development.

**Fig. 2.**
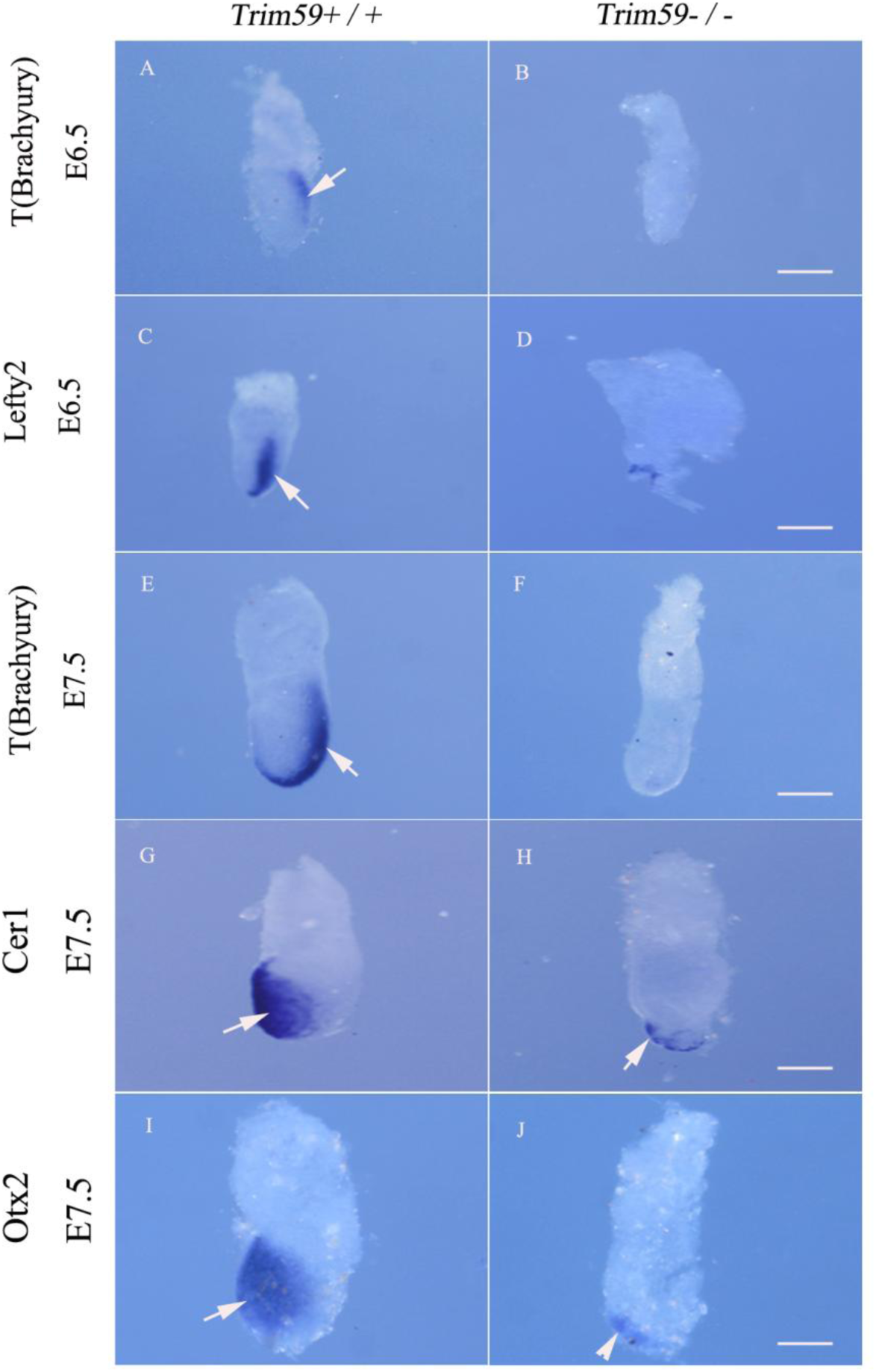
Trim59 deficiency affects expression of gastrulation-associated genes. *Trim59* +/+ and *Trim59* -/- embryos at E6.5 were hybridization using a digoxigenin-labeled anti-sense RNA probe to mouse *T (Brachyury)* (A and B) and *Lefty2* (C and D). Embryos at E7.5 were hybridization using a digoxigenin-labeled anti-sense RNA probe to mouse *T (Brachyury)* (E and F), *Cer1* (G and H) and *Otx2* (I and J). Scale bar in all panels indicates 100μm. Arrow indicates *Brachyury*, *Lefty2*, *Brachyury*, *Cer1* or *Otx2.*

PS induction depends on the crosstalk between Epi and two extra-embryonic tissues AVE and ExE. During gastrulation, signaling pathway molecules such as *Wnt3*, *BMP4*, *Nodal* and *FGF* are involved in PS formation (Padgett et al., 1993). In *Trim59* -/- embryos at E6.5, *Wnt3* expression was remarkably reduced in the EPI (Fig. 3A-C). The areas of expression *BMP4* in EPI and ExE were also smaller in size at both E6.5 and E7.5 than in wt embryos (Fig. 3D-F and G-I). Furthermore, the expression region of *BMP4* moved to EPI at E6.5 and E7.5 (Fig. 3D-F and G-I). These data show that Trim59 knockout also causes abnormal expression of signaling molecules. Taken together, Trim59 deficiency may cause the failure of gastrulation and affect the expression of gastrulation associated genes and signaling pathway molecules.

**Fig. 3.**
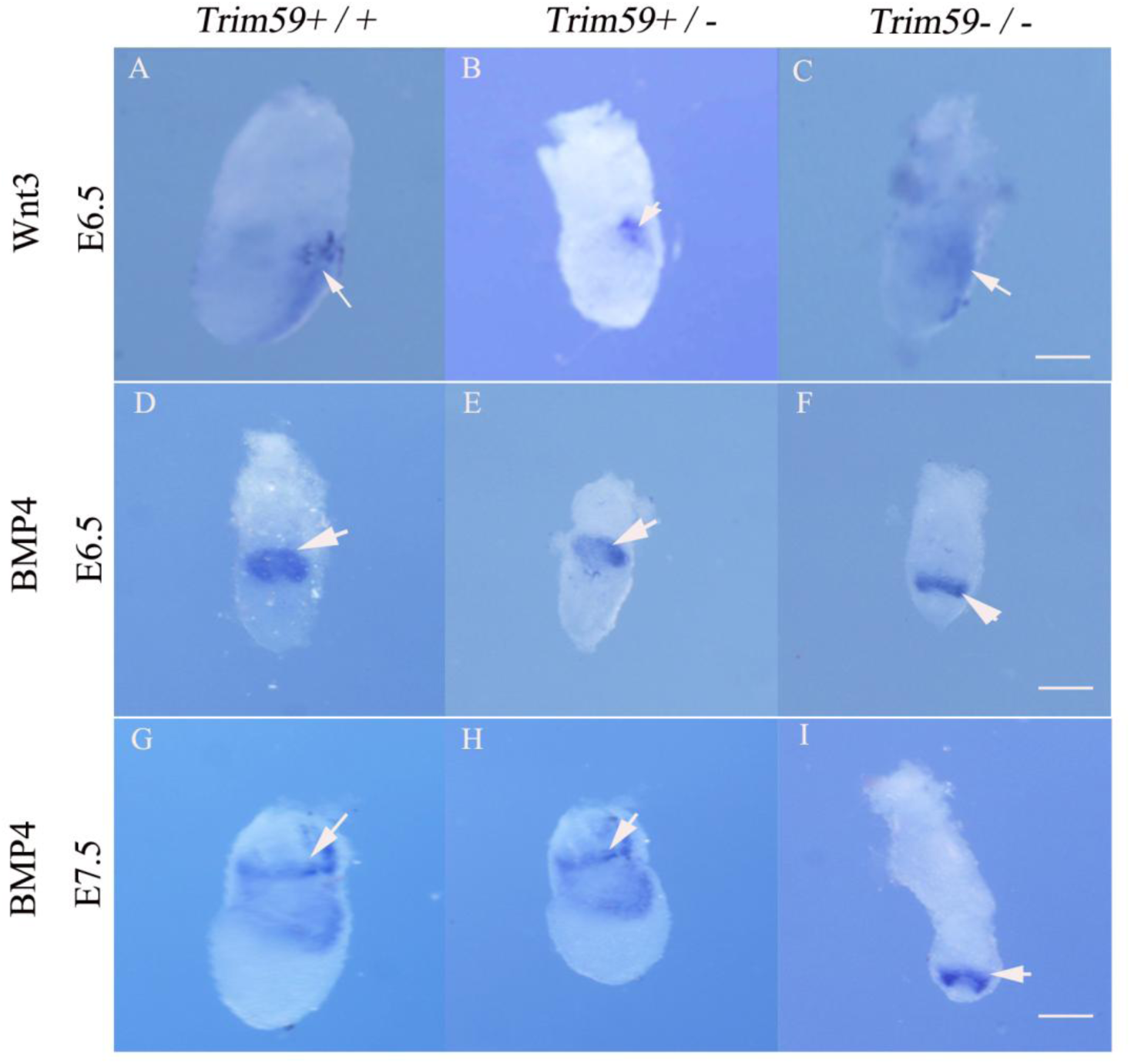
Trim59 deficiency affects Wnt3 and BMP4 expression in gastrulation stage. Embryos were hybridization using a digoxigenin-labeled anti-sense RNA probe to mouse Wnt3 (E6.5) and BMP4 (E6.5 and E7.5). Arrow indicates Wnt3 or BMP4. Scale bar, 100μm.

#### Trim59 deficiency interrupts the differentiation of blastocyst inner cell mass

The stem cells from inner cell mass (ICM) of blastocyst stage embryo are able to differentiate to generate primitive ectoderm, which ultimately differentiates into the three primary germ layers to lead the formation of gastrula (De Mot et al., 2016). Since Trim59 deficiency affects the mouse development at the gastrulation, it is possible that Trim59 deficiency may affect differentiation of ICM. To test this, we examined *in vitro* outgrowth of embryos at E3.5. At this stage, morula develops into blastocyst and EPI is formed (Sasaki, 2017). Female *Trim59* +/- mice were first superovulated and then embryos were dissected from the uterus in PBS buffer. A total of 17 outgrowths were generated. Each outgrowth was divided into two parts, one of which was used for genotyping, other blastocysts were cultured with specific stem cell medium to establish stem cell lines. PCR revealed that 12 of 17 ourgrowths were *Trim59*+/+ or *Trim59*+/- genotype, and other 5 outgrowths were *Trim59* -/- genotype (Fig. 4A and B). Importantly, while all *Trim59*+/+ 12 outgrowths grew to form stable stem cell lines, all of 5 *Trim59* -/- outgrowths failed to do this (Fig. 4A and B), indicating that Trim59 is necessary for the growth and differentiation of inner cell mass.

**Fig. 4.**
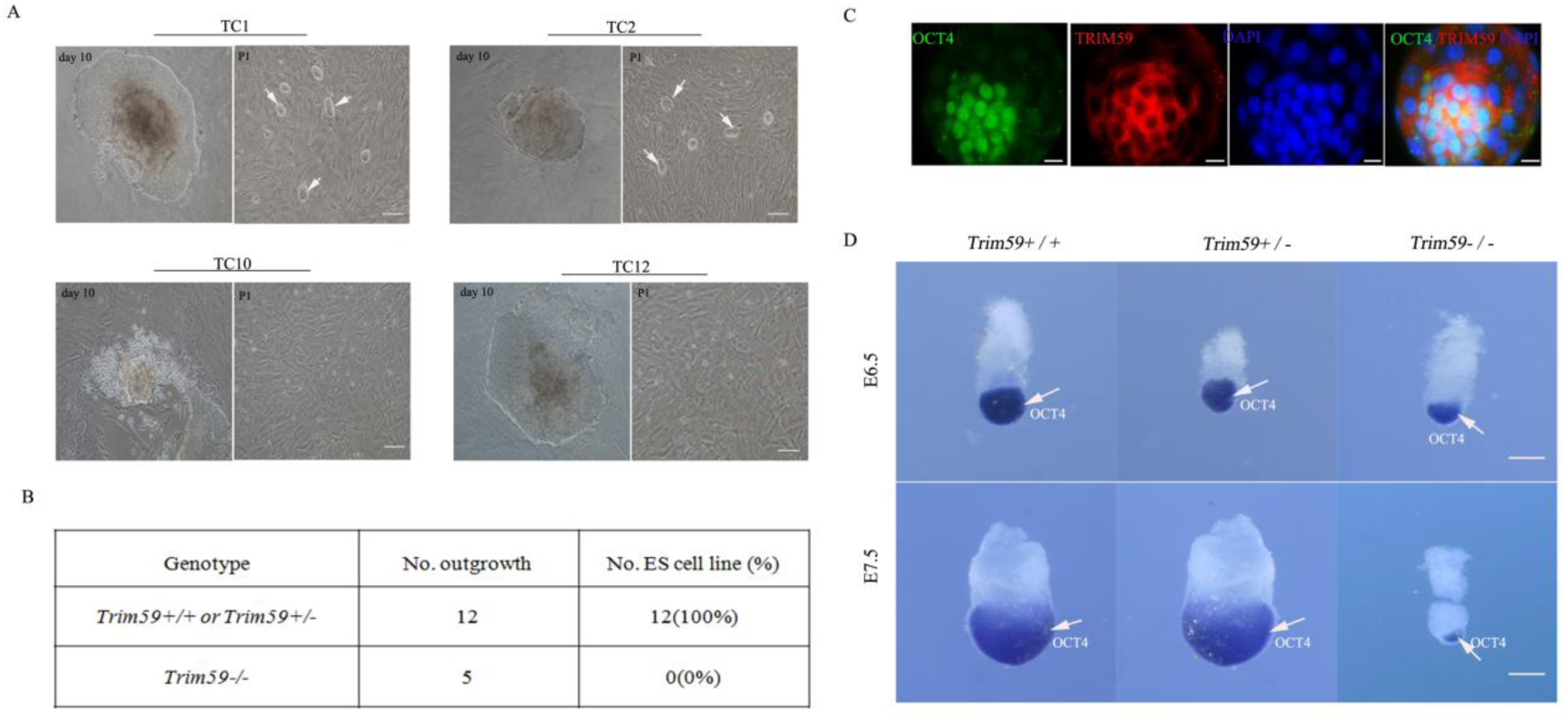
Trim59 deficiency affects differentiation of blastocyst inner cell mass. (A) ES cell clones at passage 1. TC1, TC2, TC10 and TC12, name of outgrowths. TC1 and TC2 were representative images of *Trim59* +/+ or *Trim59* +/- genotype; whereas TC10 and TC12 were representative images of *Trim59* -/- genotype. P1, passage 1. Arrows indicate ES cell clones. Scale bar, 10 μm. (B) Growth and differentiation of ES cells from *Trim59*+/+, *Trim59* +/- and *Trim59* -/- mice. No. outgrowth, number of outgrowths; No. ES cell line, number of ES cell line. (C) Staining of Trim59 and Oct4 in confluent culture of F1 ESCs. Confluent culture of F1 ESCs was stained by PE-labeled anti-mouse Trim59 and FITC-labeled anti-mouse Oct4 antibody. Images were acquired using a Bio-Rad Radiance 2100 confocal microscope with a Zeiss 63× oil immersion objective. Blue, chromatin staining. Scale bar, 10 μm. (D) Hybridization of Oct4 in *Trim59* +/+, *Trim59* +/- and *Trim59*-/- embryos. *Trim59* +/+, *Trim59* +/- and *Trim59* -/- embryos at E6.5 and E7.5 were hybridization using a digoxigenin-labeled anti-sense RNA probe to mouse Oct4. Scale bar, 100pm. Arrow indicates Oct4.

The expression of transcription factor Oct4 (octamer-binding transcription factor 4) is essential for the differentiation of blastocysts (Boiani et al., 2002; Niwa et al., 2000). Altered expression of Oct4 also marks the abnormal differentiation of blastocysts (Pesce and Scholer, 2001). Thus, we assessed the expression of Oct4 in mouse early embryos. Consistent with other data (Pfeiffer et al., 2013), Oct4 expression was restricted to ICM and EPI in wt blastocysts (Fig. 4C). Notably, during this stage, Trim59 could coexpress with Oct4 in the ICM and EPI (Fig. 4C). However, the expression region of Oct4 was remarkably decreased during gastrulation in *Trim59* -/- as compared to *Trim59* +/+ embryos (Fig. 4D), further indicating that ICM differentiation is abnormal in *Trim59* -/- blastocyst stage embryos. Taken together, our data suggest that Trim59 is a critical factor for ICM differentiation in blastocyst stage embryos.

#### Trim59 deficiency disturbs F-actin polymerization during ICM differentiation

Since Trim59 plays a critical role in blastocyst development stage, next question asked is how Trim59 regulates the formation of blastocysts. We first employed yeast two-hybrid method to found potential target molecules of Trim59. Trim59 potentially interacted with multiple proteins (only eight proteins were listed) (Table 2), some of which are relative to the function of cytoskeleton in early embryo, including actinin alpha 1(ACTN1), phospholipid scramblase 1 (PLSCR1) and F-actin binding protein (TRIOBP). Immunoprecipitation-Mass Spectrometry further revealed that Trim59 interacted with myosin and F-actin capping protein (Fig. S3), which are also associated with the cellular cytoskeleton functions during early embryo development. Deficiency or dysregulation of these genes involved in the regulation of actin skeleton may lead to impaired embryonic development and lethality (Boumela et al., 2011; Ke et al., 2010; Lanier et al., 1999; Niedenberger et al., 2014). So we proposed that the effects of Trim59 on blastocyst development may be through modulating actin cytoskeleton. To test this hypothesis, we employed a loss-of-function experiment. In untreated F1 ESCs, F-actin was polymerized at the cell-cell junction and at the edge of the cultivated limbal stem cell (Fig. 5A). However, while F1 ESCs were treated by silencing Trim59, F-actin polymerization at the cell-cell junction and at the edge of the cultivated limbal stem cell was interrupted (Fig. 5A). We then examined blastocysts of *Trim59* -/- embryos. At E3.5 and E4.5, the blastocysts in *Trim59*-/- embryos had weaken assembly of F-actin at the edge of cell-cell junction as compared to wt embryos (Fig. 5B). Trim59 contains three functional domains: a RING-finger, a coiled-coil region, and a B-box domain. We next further determined which domain of Trim59 induces polymerization of F-actin. FLAG-tagged Trim59 expression vector and four different deletion mutants of Trim59 were generated (Fig. 6A-B). Only Trim59 full-length (T1) and Trim59 fragments (T4 and T5) which contain a RING-finger domain could promote the assembly of F-actin at the boundary of cells and maintain cells in the stretched state; Whereas in Trim59 fragments (T2 and T3) transfected HEK293T cells, which are absence of RING-finger domain, there had a decreased immunofluorescence intensity, rounded cells edge, degraded diopter and concomitant morphological changes (Fig. 6C and Fig. S5) as compared to cells transfected by fragments containing RING-finger domain. Thus, RING-finger domain of Trim59 is necessary for the regulation of F-actin assembly. Taken together, the effects of Trim59 on inner cell mass differentiation are through disturbing F-actin polymerization.

**Fig. 5.**
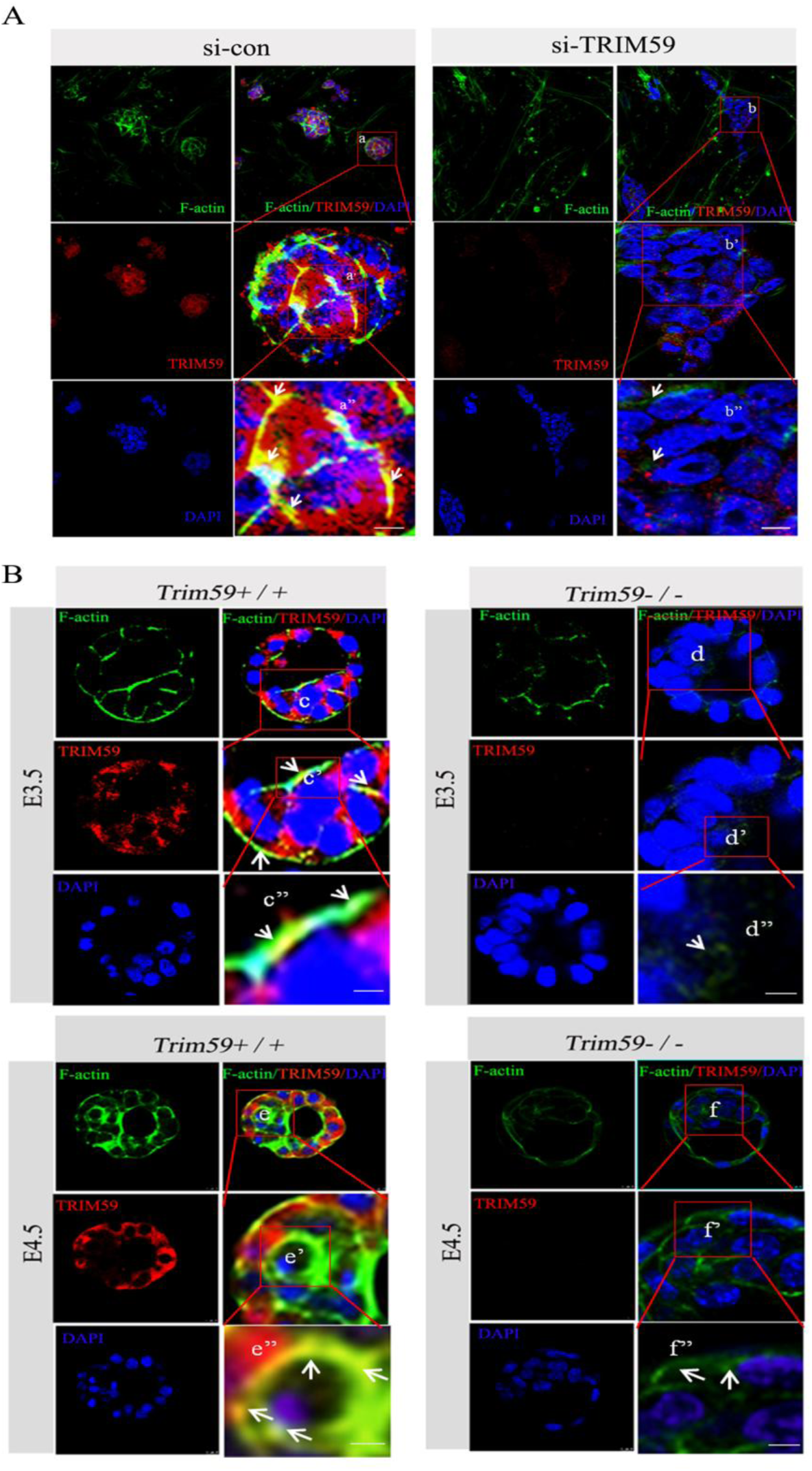
Trim59 deficiency affects F-actin polymerization. (A) Immunofluorescence assay of Trim59 and F-actin in F1 ESCs transfected with Trim59 siRNA (si-TRIM59) or siRNA control (si-con). F-actin (green), Trim59 (red) and chromatin (blue). Images were acquired using a Bio-Rad Radiance 2100 confocal microscope with a Zeiss 63× oil immersion objective. (B) Confocal fluorescence images of *Trim59* +/+ or *Trim59* -/- embryos at E3.5 and E4.5 stained with Alexa488, F-actin (green), Trim59 (red) and DAPI (blue). Images were acquired using a Bio-Rad Radiance 2100 confocal microscope with a Zeiss 63× oil immersion objective. Scale bar, 10 μm; Arrows indicate F-actin polymerization.

**Fig. 6.**
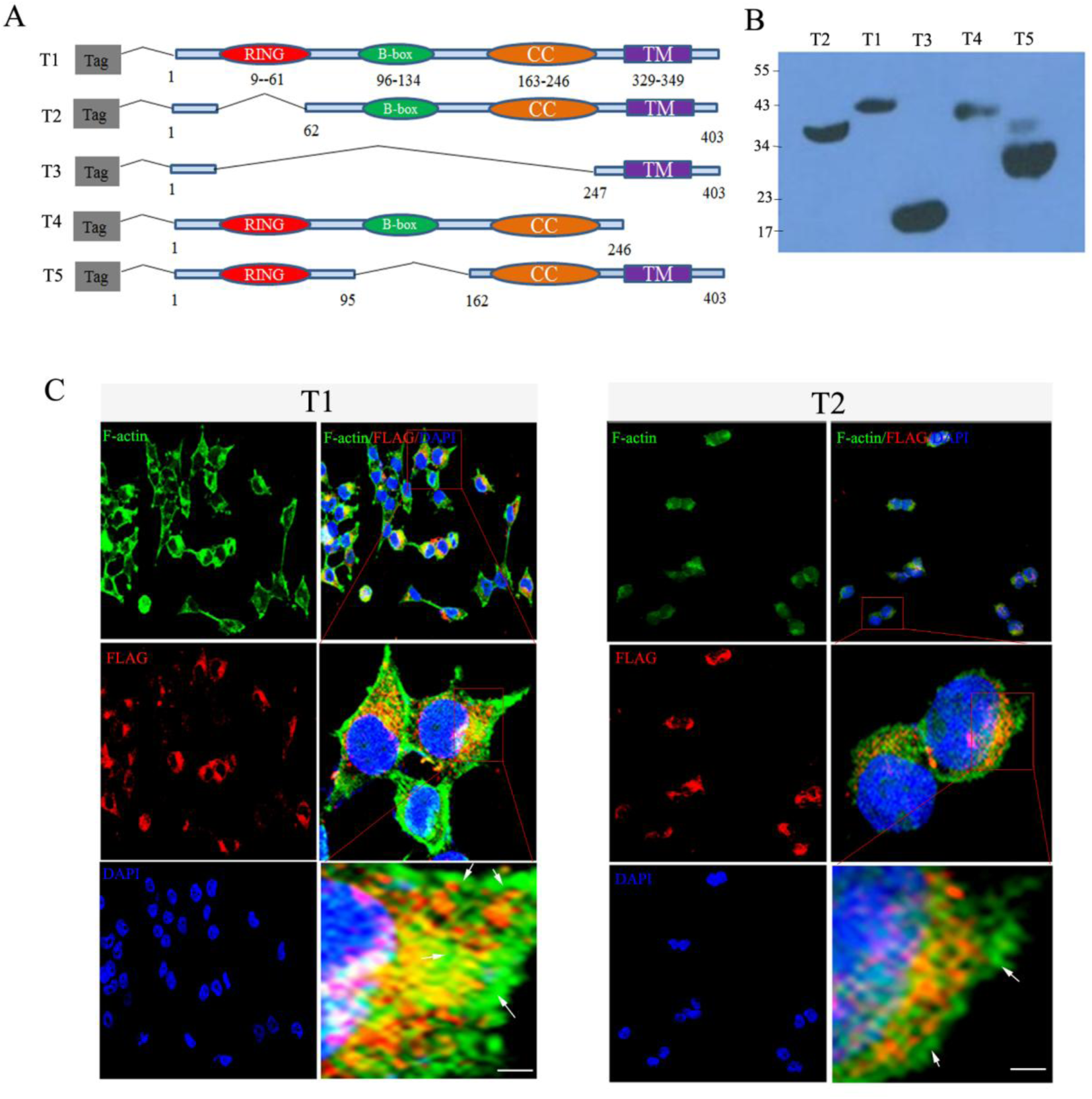
Trim59 RING-finger domain is required for F-actin assembly. (A) Schematic representation of full-length Trim59 and its domain deletion mutants. RING, RING-finger domain; B-box, B-box domain; CC, coiled-coil domain; TM, transmembrane domain. T1, full-length Trim59; T2, RING-finger domain deleted fragment; T3, RING-finger domain, B-Box domain and coiled-coil domains deleted fragment; T4, C terminal TM deleted fragment; T5, B-box deleted fragment. (B) Immunoblotting of Trim59 and its fragments. FLAG-tagged plasmids were constructed and transfected into HEK293T cells for 24hrs. The expression of Trim59 full-length (T1) and different domain mutants (T2-T5) was analyzed by anti-FLAG primary antibody. (C) Immunofluorescence assay of F-actin in FLAG-tag full-length Trim59 or its fragment transfected HEK293T cells. HEK293T cells were transfected with vectors encoding FLAG-full length Trim59 (T1) or its FLAG–tagged RING-finger domain deleted fragment (T2) for 24hrs. F-actin, green; Trim59, red; Chromatin, blue. Scale bar, 10 μm.

**Table 2.** Proteins that potentially interact with Trim59 (data from yeast two-hybrid).

| Name | Accession number | Description |
| --- | --- | --- |
| ACTN1 | NM_001102 | *Mus musculus* actinin, alpha 1 |
| ARNT | NM_178427 | *Mus musculus* aryl hydrocarbon receptor nuclear translocator |
| CD74 | NM_004355 | *Mus musculus* CD74 antigen |
| HOMER3 | NM_004838 | *Mus musculus* homer homolog 3 |
| PLSCR1 | NM_021105 | *Mus musculus* phospholipid scramblase 1 |
| RPS2 | NM_002952 | *Mus musculus* ribosomal protein S2 |
| TRIOBP | NM_007032 | *Mus musculus* TRIO and F-actin binding protein |

#### Trim59 promotes polymerization of F-actin via WASH K63-linked polyubiquitination

Previous studies have shown that TRIM proteins with RING-finger domain may act as an E3 ubiquitin ligase (Esposito et al., 2017; Li et al., 2014; Yan et al., 2014). Next, we investigated whether Trim59 mediated F-actin polymerization is through its ubiquitin ligase activity. We first examined which protein was involved in Trim59 mediated F-actin polymerization. WASH, a member of Wiskott–Aldrich syndrome protein (WASP) family, plays an important role in regulating polymerization of F-actin (Fig.7A) (Wang et al., 2014; Xia et al., 2013). Thus we hypothesized that Trim59 could promote WASH ubiqutination through ubiqutin-protease system. Because ubiqiotin linkages via lysine 48 (K48) or 63(K63) can differentially address proteins for 26S proteasomal degradation or a common activating signal (Deng et al., 2000), we transfected HEK293T cells to express WASH and Trim59 in the presence of vectors encoding K48-linked ubiquitin or K63-linked ubiquitin. Trim59 resulted in more K63-linked ubiquitin but not K48-linked ubiquitin on WASH protein (Fig. 7B). Meanwhile we also utilized a K220R WASH mutant, which is an ubiquitin variant and unable to produce a specific ubiquitin chain (Hao et al., 2013) as a control. We found that Trim59 did not increase WASH K220R mutant K63-linked ubiquitin (Fig. 7C). RING-finger domain deletion mutants (T2 and T3) failed to promote WASH K63-linked ubiquitin (Fig. 7D). We also immunoprecipitated endogenous WASH and measured total Ub, K63-linked or K48-linked ubiquitin associated with WASH by silencing expression of Trim59 with specific siRNA. There was remarkable total Ub and K63-linked ubiquitin; Whereas total Ub and K63-linked ubiquitin were obviously decreased on the WASH protein after Trim59 was silenced (Fig. 7E). No K48-linked ubiquitin was found in both control and Trim59 siRNA transfection groups (Fig. S7). These data suggest that WASH may be involved in Trim59 mediated assembly of F-actin. Indeed, when the expression of WASH protein was interrupted by specific siRNA *in vitro*, stem cell mass was irregular than those by control transfection (Fig. 7F). Especially some smaller cell masses appeared after silencing WASH (Fig. 7F). Meanwhile, F-actin polymerization decreased at the cytomembrane and cell-cell junction (Fig. 7F) although stem cell morphology was not changed. These results suggest that Trim59 - mediated assembly of F-actin is dependent on WASH K63-linked ubiquitination.

**Fig. 7.**
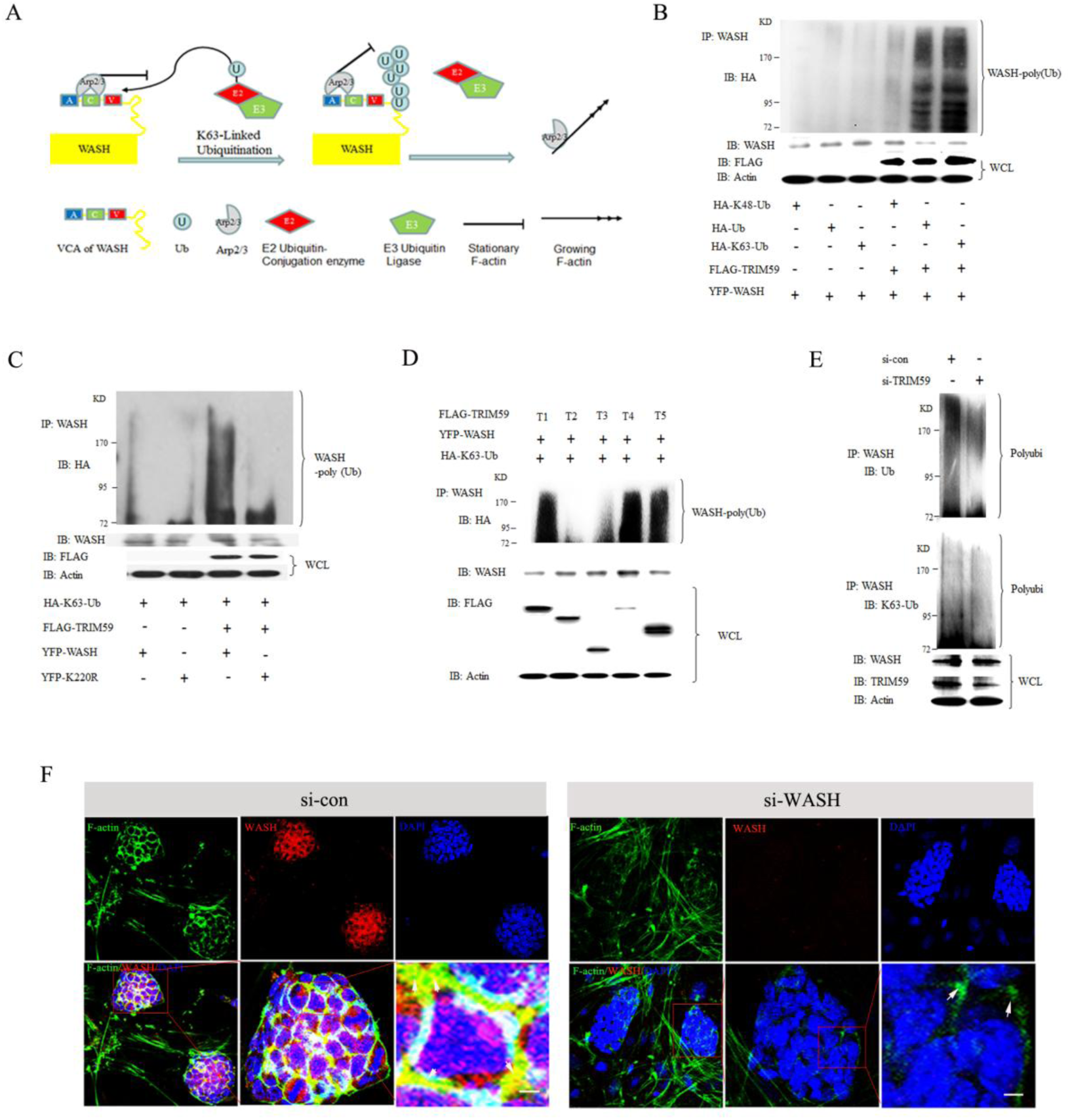
Trim59 promotes WASH K63-linked ubiquitination. (A) Model of WASH, Arp2/3, E2, E3 and K63 ubiquitin regulation on F-actin polymerization. (B) Immunoblotting of WASH K63-linked ubiquitin in cotransfected HEK293T cells. HEK293T cells were cotransfected with YFP-WASH and with (+) or without (-) FLAG-Trim59 as well as with HA-Ub, HA-K48-Ub or HA-K63-Ub. Immunoprecipitation was performed by anti-WASH. Trim59 and WASH were detected by anti-FLAG and anti-WASH antibodies. The polyubiquitination of WASH was detected by anti-HA. IP, immunoprecipitation; WCL, whole cell lysate; Actin, a loading control. (C) Immunoblotting of WASH K63-linked ubiquitin in cotransfected HEK293T cells. HEK293T cells were cotransfected with YFP-WASH or YFP-K220R and with (+) or without (-) FLAG-Trim59 as well as with HA-K63-Ub. Immunoprecipitation was performed by anti-WASH. Trim59, WASH and WASH variant were detected by anti-FLAG and anti-WASH antibodies. Polyubiquitination of WASH was detected by anti-HA. IB, immunoblot assay; IP, immunoprecipitation; WCL, whole cell lysate. (D) Immunoblotting of WASH K63-linked ubiquitin in cotransfected HEK293T cells. HEK293T cells were cotransfected with FLAG-Trim59 full length (T1) and different domain deletion mutants (T2 toT5) and with YFP-WASH (+) as well as with HA-K63-Ub (+). Immunoprecipitation was performed by anti-WASH. Trim59, Trim59 fragments and WASH were detected by anti-FLAG and anti-WASH antibodies. Polyubiquitination of WASH was detected by anti-HA. IB, immunoblot assay; IP, immunoprecipitation; WCL, whole cell lysates; Actin, a loading control. (E) Immunoblotting of K63-linked ubiquitination (K63-Ub) of endogenous WASH from F1 ESCs after treated with specific Trim59 siRNA. Immunoprecipitation was performed by anti-WASH. Trim59 and WASH were detected by anti-Trim59 and anti-WASH antibodies. Immunoblot analysis of Ub-WASH (top blot), K63-Ub-WASH (below blot) by anti-Ub and anti-K63 antibodies. Polyubi., polyubiquitination; IP, immunoprecipitation; IB, immunoblot assay; Actin, a loading control; WCL, whole cell lysates. (F) Immunofluorescence assay of F-actin assembly in WASH siRNA or siRNA control transfected F1 ESCs. Mouse F1 ESCs was transfected with siRNA control (si-Con) or mouse WASH siRNA (si-WASH) for 72 hrs. F-actin, green; WASH, red; chromatin, blue. Scale bar, 10 μm.

## DISCUSSION

In this study we found that Trim59 is a critical regulator for early embryo development from blastocyst stage to gastrula through modulating F-actin assembly. *Trim59* -/- mouse embryos are failure for normal embryogenesis and have a reduced expression of primary germ layer formation associated genes including *Brachyury*, *lefty2*, *Cer1*, *Otx2*, *Wnt3* and *BMP4*. We found that Trim59 deficiency may disturb F-actin polymerization during inner cell mass differentiation. We also demonstrate that the effects of Trim59 on F-actin polymerization is through WASH K63-linked ubiquitination. Overall, these results provide a molecular basis for early embryonic development. More broadly, our findings are relevant to understand the impact of Trim59 on human infertility and embryonic lethality.

TRIM family members have been implicated in a variety of biological processes, such as the regulation of differentiation and development (James et al., 2007; Kitamura et al., 2005). Several members of TRIM proteins have been found to play a crucial role during early embryonic development. TRIM33 regulates ectodermal induction by functioning as a smad4 ubiquitin ligase (Kim and Kaartinen, 2008). TRIM71 has RING-dependent ubiquitin ligase activity. *Trim71*-/- embryos present a highly penetrant closure defect of the cranial neural tube, and cease development and die between E9.5 and E11.5 (Cuevas et al., 2015). TRIM36 also markedly and specifically inhibits somite formation and vegetal microtubule polymerization (Houston and Cuykendall, 2009). Trim36-depleted embryos are disrupted in the development of cortical rotation in a manner dependent on ubiquitin ligase activity. We here found that Trim59 plays a critical role in early embryos development from blastocyst stage into gastrula. Trim59 knockout affects primary germ layers formation at the beginning of gastrulation. Thus, multiple members of TRIM family may be involved in early embryos development.

Our studies suggest that Trim59 is necessary for the formation of primary germ layers, which happens at the beginning of gastrulation. Transition from blastocyst to gastrula is a remarkably elaborate process involving a multiple of genes such as *Brachyury*, *Otx2*, *Cer1* and *Lefty2* (Parfitt and Shen, 2014). *Brachyury* is a key player in mesoderm formation (Thomson et al., 2011). *Brachyury* defection fails to elongate along the anterior-posterior axis and their embryos can not develop mesodermal extraembryonic tissues (Herrmann, 1991). Our data showed that *Brachyury* is not detected in *Trim59* -/- embryonic blastopore, suggesting Trim59 deficiency affects the mesoderm formation. *Otx2*, *Cer1* and *Lefty2* belong to AVE genes (Goncalves et al., 2011a), expression of these genes could not also be detected in Trim59 deficient embryos, indicating that AVE formation may be regulated by Trim59. Many signaling molecules participate in regulating early embryos development such as BMP (Basilicata et al., 2016; Kurek et al., 2015), Nodal (Spiller et al., 2012), Wnt (Barrow et al., 2007), and FGF (Wilcockson et al., 2017). Wnt3 activity derived from the posterior visceral endoderm has a temporal role in establishing the primitive streak (Yoon et al., 2015). Wnt3 co-receptor Lrp6 knockout embryos fail to establish a primitive streak (Kelly et al., 2004). Extra-embryonic ectoderm (ExE) expresses BMP4, which is involved in many steps in pregastrulation development. The loss of BMP4 functions leads to gastrulation defects (Blitz et al., 2003). In *Trim59* -/- embryos, there have a remarkably reduced expression in both Wnt3 and BMP4. The region of expression BMP4 also moves to the EPI. Oct4 is expressed in mouse totipotent embryonic stem and germ cells, and totipotent cells differentiate into somatic lineages occurred at the blastocyst stage and during gastrulation (Le Bin et al., 2014). When Oct4 is deleted, embryonic stem (ES) cells lose the capacity to self-renew and subsequently differentiate into extra-embryonic trophectoderm (Niwa et al., 2000). *Oct4*-/- embryos die at peri-implantation stages due to the conversion of ICM into trophectoderm (Nichols et al., 1998). At E6.5 and E7.5, Trim59 knockout also decreases the expression of Oct4.

Our mechanistic studies uncovered that K63 linked ubiquitination of WASH by Trim59 is required for polymerization of F-actin during gastrulation. Actin microfilaments are the major regulators and play a crucial role in cell morphology and mobility (Montazeri et al., 2015). We found that Trim59, as an E3 ligase family member, promotes WASH K63-linked polyubiquitination through RING finger domain. Recent studies have also indicated that regulation of WASH-dependent actin polymerization is based on K63 ubiquitination in WASH (Hao et al., 2013). WASH is a member of the Wiskott-Aldrich syndrome protein family consisting of WASP/N-WASP, WAVE, WHAMM, JMY, and WASH (Campellone and Welch, 2010). WASH contains a carboxyterminal VCA (verprolin homologous or WH2, central hydrophobic, and acidic) motif that binds to actin and Arp2/3 complex to stimulate actin filament nucleation (Derivery et al., 2009; Jia et al., 2010). Ubiquitination is a posttranslational modification which can have pleiotropic effects on its substrates depending on the length and type of ubiquitin chains. Studies have shown that K63 linked ubiquitination typically acts as a signaling event to modify function, such as altering protein-protein interactions, protein conformations, or targeting proteins for lysosomal delivery (Sun and Chen, 2004). Our data exhibit that WASH K63-linked ubiquitination by Trim59 determines the polymerization of F-actin during gastrulation.

## MATERIALS AND METHODS

### Generation of Trim59 knockout mice

In brief, Trim59 gene was retrieved from a 129/sv BAC clone bMQ452g13 (provided by Sanger Institute) by a retrieval vector containing two homologous arms. After correct recombination, this vector contains 11.3kb of genomic sequence including part of intron II, exon III and 3.5kb downstream sequences. 147bp intron II and the entire coding region of Trim59 were then deleted and replaced with a loxP-Neo-loxP cassette. The targeting construct which contains a neo cassette for positive selection and a herpes simplex virus-thymidine kinase expression cassette for negative selection was linearized with *Not* I and electroporated into C57BL/6 derived B6/BLU embryonic stem (ES) cells. 96 ES cell clones were selected and verified for correct recombination with long range PCR and Southern blot analysis. Correctly targeted ES cells were injected into C57BL/6J blastocysts followed by transfer to pseudopregnant mice. Chimerical male mice identified by PCR were bred to C57BL/6J females to generate F1 offspring. Germ line transmission of the targeted Trim59 allele was verified by PCR analysis of tail DNA from F1 offspring with agouti coat color. All procedures were conducted in accordance with Institutional Animal Care and Use Committee of Model Animal Research Center. Primers used in this study were listed in supplementary Table S3A.

### Reagents and plasmid constructs

Mouse Trim59 siRNA, WASH siRNA and control mock siRNA were purchased from Ribo, Shenzheng, China. Following antibodies were purchased including anti-Trim59 (Abcam), anti-WASH (Sigma), anti-Oct4 (Abcam), anti-FLAG (Abmart), anti-HA (Cell signaling technology), anti-Ub (Immunoway), anti-K48 (Cell signaling technology), anti-K63(Enzo) and anti-β-actin (Santa). Phalloidin-iFluor 488 (Abcam), DAPI (Cell signaling technology), Alexa-488 and Alexa 594 (Cell signaling technology) were also purchased.

Murine full-length Trim59 clone was obtained from the ATCC. Trim59 mutants were constructed by performing PCR with four primers according to a previous method (Zhang et al., 2009). The PCR products were then ligated into a p-FLAG-CMV™-1 mammalian expression vector (Invitrogen). All the constructs were confirmed by DNA sequencing (Huada Bio., China). All primers used in this study are listed in supplementary Table S3B. YFP–WASH and YFP-WASH K220R mutant were from Patrick Ryan Potts, UT Southwestern Dallas, TX 75390, USA; Plasmids encoding hemagglutinin (HA)-tagged ubiquitin (HA-Ub), hemagglutinin-tagged K48-linked ubiquitin (HA-K48-Ub) and K63-linked ubiquitin (HA-K63-Ub) were obtained from Y. Xiong (University of North Carolina, Chapel Hill).

### Embryos dissection and genotyping

*Trim59* +/- mice were bred to obtain wild-type (*Trim59* +/+), heterozygote (*Trim59* +/-), and homozygous mutant (*Trim59* -/-) embryos. The female mice were first superovulated by intraperitoneal injection of 5 IU pregnant mare serum gonadotropin (PMSG), and followed by injection of 5 IU human chorionic gonadotropin (hCG) 48hrs later, and then mated with male *Trim59* +/- mice. Females were screened for vaginal plugs following morning (E0.5). Embryos at E3.5, E4.5, E6.5, E7.5, E8.5 and E9.5 were collected from the uterus in PBS buffer, and part of embryo tissue was used for genotyping.

### RT-PCR and qRT-PCR

Semi-quantitative reverse transcription-polymerase chain reaction (RT-PCR) and quantitative real-time PCR (qRT-PCR) were performed. Briefly, total RNA was extracted from the cells, tissues and organs using TRIzol reagent (Invitrogen Corp). First-strand cDNA was generated from the total RNA using oligo-dT primers and reverse transcriptase (Invitrogen Corp). The PCR products were visualized on 1.0% (wt/vol) agarose gel. Real-time PCR was conducted using QuantiTect SYBR Green PCR Master Mix (Qiagen) and specific primers in an ABI Prism 7000 analyzer (Applied Biosystems). GAPDH mRNA expression was detected using each experimental sample as an endogenous control. The fold changes were calculated using the ΔΔC_t_ method according to the manufacturer’s instructions (Applied Biosystems). All the reactions were run in triplicate. Primer sequences are listed in Supplementary Table S3C and D.

### RNA interference

F1 ESCs were transfected with murine Trim59 siRNA, WASH siRNA or mock siRNA by Hiperfect Transfection Reagent (siRNA transfection) (Qiagen, Valencia, CA, USA). Interference sequences were designed according to pSUPER system instructions (Oligoengine). After transfection for 72 hrs, the cells which cultured at slide were washed with PBS for twice for immunofluorescent analyses. Other parts of the cells were harvested for RNA extraction for qRT-PCR and immunoblot analyses to test the interference efficiency. siRNA sequences are listed in Supplementary Table S3E.

### Immunoblot, immunoprecipitation and liquid chromatography-tandem Mass

For immunoprecipitation, HEK293T cells were transfected with YFP -WASH, YFP-WASH K220R mutant, FLAG-Trim59, HA-Ub, HA-K48 Ub and/ or HA-K63 Ub vectors for 24hrs. The cells were lysed with cell lysis buffer (Cell Signaling Technology), which was supplemented with a protease inhibitor ‘cocktail’ (Calbiochem). The protein concentrations of the extracts were measured using a bicinchoninic acid assay (Pierce). Immunoprecipitation (IP) was performed as described by the manufacturer (Thermo Scientific, USA). Cell lysates were preabsorbed with protein A/G agarose beads at 4°C for 1 h. After centrifuging, the supernatants were incubated with anti-WASH antibody overnight at 4°C followed by incubation with protein A/G agarose beads (Santa Cruz biotechnology) for 2 hr at 4°C. The precipitates were collected and completely washed with lysis buffer for 3 to 5 times. After centrifuging, supernatants were cultured with anti-WASH (1:1000) and anti-HA (1:5000) at 4°C for overnight, and the secondary antibodies (Boehringer Mannheim) and enhanced chemiluminescence (Amersham). The gel lanes containing the immunopurified samples were excised for Liquid Chromatography-tandem Mass (LC-MS/MS) analysis.

For immunoblot, hybridizations with primary antibodies were conducted for 1 h at room temperature in blocking buffer. The protein-antibody complexes were detected using peroxidase-conjugated secondary antibodies (Boehringer Mannheim) and enhanced chemiluminescence (Amersham).

### Histology and immunofluorescence

For hematoxylin/eosin (H&E) staining of embryos, *Trim59*+/+ and *Trim59* -/- embryos at E6.5 and E7.5 were fixed in 10% formalin-buffered saline and embedded in paraffin, 5 μm sections kidney sections were cut and stained with H&E.

For immunofluorescence staining of embryos, E3.5 and E4.5 embryos collected from uterus were washed with PBS for 3 times and then fixed for 30 min in 4% PFA. Fixed embryos were permeabilized in 0.1% Triton X-100 in PBS for 30 min. 5% goat serum was used for blocking. Primary anti-Trim59 (1:500) was added for 2hrs. After washing with PBS the coverslips were stained with phalloidin (1:200) and Alexa 488 conjugated secondary antibodies for 1hr. After washing with PBS for 3 times, nucleus was stained with DAPI for 3 min. Embryos were placed in glycerin droplet for fluorescence microscopy immediately. After that embryos were digested to extract genome DNA for genotyping PCR.

For immunofluorescence staining of cells, F1 ESCs were cultured on slides and transfected by siRNA for 72hrs. ES cells were washed with PBS for 3 times, fixed for 20 min in PBS containing 4% PFA at room temperature, and then permeabilized in 0.1% Triton X-100 in PBS for 30 min. 5% goat serum was used for blocking. Primary anti-Trim59 (1:500) or anti-WASH (1:500) antibodies were added for 2hrs. After washing with PBS, the coverslips were stained with phalloidin (1:200) and Alexa 488 conjugated secondary antibodies for 1hr. Immunofluorescence images were obtained with laser scanning confocal microscopy.

### Mouse embryo stem cell culture

Intact blastocysts from *Trim59* +/- crosses were seeded on the feeder layers of mitomycin C-treated MEF in the ES medium consisting of knockout DMEM (Gibco), 20% KSR, supplemented with 1000 units / mL mouse leukemia inhibitory factor (LIF) (ESGRO, Chemicon), 0.1 mM non-essential amino acid (NEAA), 1 mM L-glutamine, 0.1 mM β-mercaptoethanol, penicillin (100μg/ml) and streptomycin (100μg/ml). Half of the medium was changed daily. Approximately 10 days after seeding, inner cell mass outgrowth was mechanically removed and divided into two parts: one part was collected for genotyping and the other was digested with 0.25% trypsin–EDTA into small clumps for ES cell line derivation. Digestion was halted with trypsin inhibitor and the outgrowths reseeded on fresh feeder cells. All ES cell lines then were massaged and cultured in 20% FBS (Hyclone) ES medium (instead of KSR ES medium). ES cell lines were stored in freezing medium (ES medium supplemented with 10% DMSO and 40% FBS), and frozen in liquid nitrogen.

### *In situ* hybridization

Timed pregnancies between *Trim59* +/- male and *Trim59* +/- female mice were established and vaginal plug identification was designated as embryonic day 0.5. Embryos (E4.5-E10.5) were harvested through dissected from uterus then embryos were dissected into PBS, amniotic and yolk sac were used to identify embryonic genotyping. Embryos (E6.5 and E7.5) were fixed in 4% PFA for 3hrs, washed in PBS, and dehydrated in ethanol at an increasing series of concentrations from 70% to 100%. Then embryos were rinsed in methylsalicylate and embedded in paraffin. The samples were sectioned into 5μm slides and then hybridized with corresponding probes. Digoxigenin-labeled anti-sense RNA probes to mouse *Oct4*, *Brachyury*, *Bmp4*, *Wnt3*, *Cer1*, *Lefty2*, *Otx2* and *Trim59* gene were used *in situ* hybridization. The sense probes were used as negative controls. The sequence of associated gene probes were listed in supplementary Table S3F.

### *Yeast* two-hybrid analyses

Yeast two-hybrid analysis was performed by Beijing Proteome Research Center. The ProQuest yeast two-hybrid system (Y2H) (Invitrogen) was employed to assess interactions between Trim59 and other subfamily members. Trim59 was first cloned into pDBleu as a bait plasmid pDBleu-Trim59. A mouse embryos cDNA library fused to GAL4AD of pEXP-AD502 (Invitrogen) was screened for proteins that interact with Trim59 using the ProQuest Two-Hybrid System (Invitrogen). The detailed method of yeast two-hybrid screening has been described previously (Invitrogen). Briefly, the pEXP-AD502 plasmid and the pDELeu-Trim59 bait plasmid were co-transformed into AH109 yeast cells. Transformed yeast cells were plated on medium lacking histidine or uracil or medium containing 5-fluoroorotic acid (5FOA). The transformed yeast cells were also plated on YPAD plates to further conduct β-galactosidase assays. A total of 1×10e7 library clones were screened for growth on selective media and assayed for β-galactosidase activity. pEXP-AD502 cDNA plasmids were recovered by bacterial transformation of DNA isolated from positive yeast colonies. The candidate pEXP-AD502 cDNA plasmids were retransformed into yeast cells with the empty pDEST32 vector or pDELeu-Trim59 plasmid encoding irrelevant bait to exclude false-positives. Inserts of true-positive pEXP-AD502 cDNA clones were characterized by sequence analysis.

### Statistical analyses

All quantitative data were expressed as mean ± SEM. Significance was evaluated with a two-tailed unpaired Student’s t-test. Excel and Prism Version 5 software (GraphPad) were used for statistical evaluation. A 95% confidence interval was considered significant and was defined as *p*<0.05.

## Acknowledgements

We are grateful to the members of Lin Liu Lab and the anatomy institute of Shandong university for assistance and discussion.

## Competing interests

The authors declare no competing or financial interests.

## Author Contributions

R.Y. designed research and wrote paper; X. S. designed research, conducted experiments (mechanism) and wrote paper; C. W. and Y. L. designed and conducted experiments (embryonic observation). X. Y. and L. L. designed and conducted experiments (embryo stem cell). M. Z., S. M., S. W. were involved in experiments; Z. Z. participated in study design; Y. C. and Y. Z. offered assistances for the animal experiments. All authors read and approved the final manuscript.

## Funding

This research was supported by NSFC grants 91029736, 9162910, 81600436 and 91442111, the Joint NSFC-ISF Research Program, which is jointly funded by the National Natural Science Foundation of China and the Israel Science Foundation (ISF-NSFC program 31461143010), a Ministry of Science and Technology grant (863 program, 2008AA02Z129), and national key research and development program of china (2016YFC1303604); the Program for Changjiang Scholars and Innovative Research *Team* in University (No. IRT13023) and the State Key Laboratory of Medicinal Chemical Biology.

## Supplementary data

**Fig. S1.**
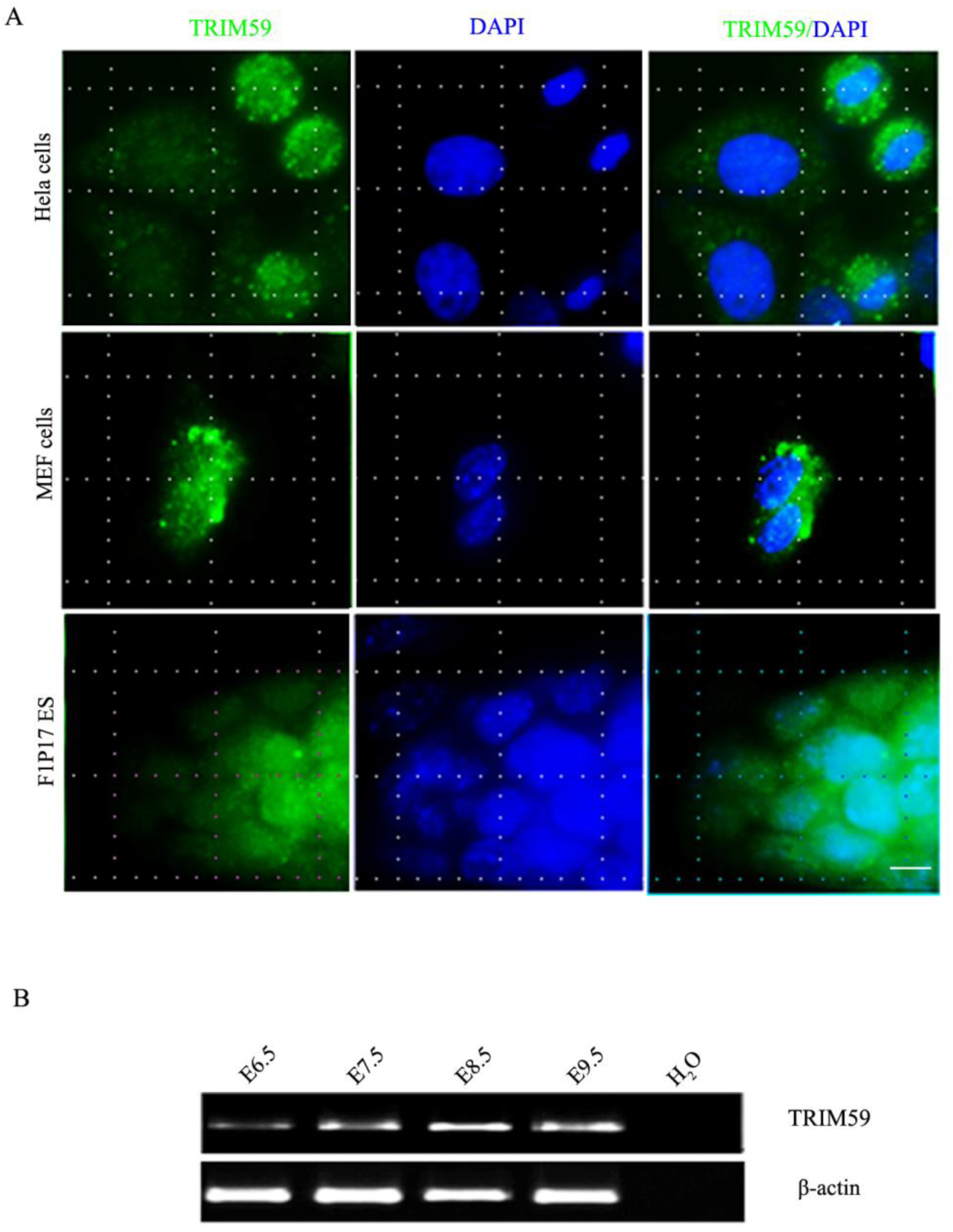
Expression of Trim59 in different cell lines and mouse early embryos. **(A)** Immunofluorescence assay of Trim59 in Hela cells (human cervical cancer cell), MEF (murine Embryonic Fibroblast) cells and F1 ESCs. Trim59 (green) and chromatin (blue). Images were acquired using a Bio-Rad Radiance 2100 confocal microscope with a Zeiss 63× oil immersion objective. Scale bar, 10 μm. (B) RT-PCR of Trim59 in mouse early embryos from E6.5 to E9.5. β-actin, a loading control.

**Fig. S2.**
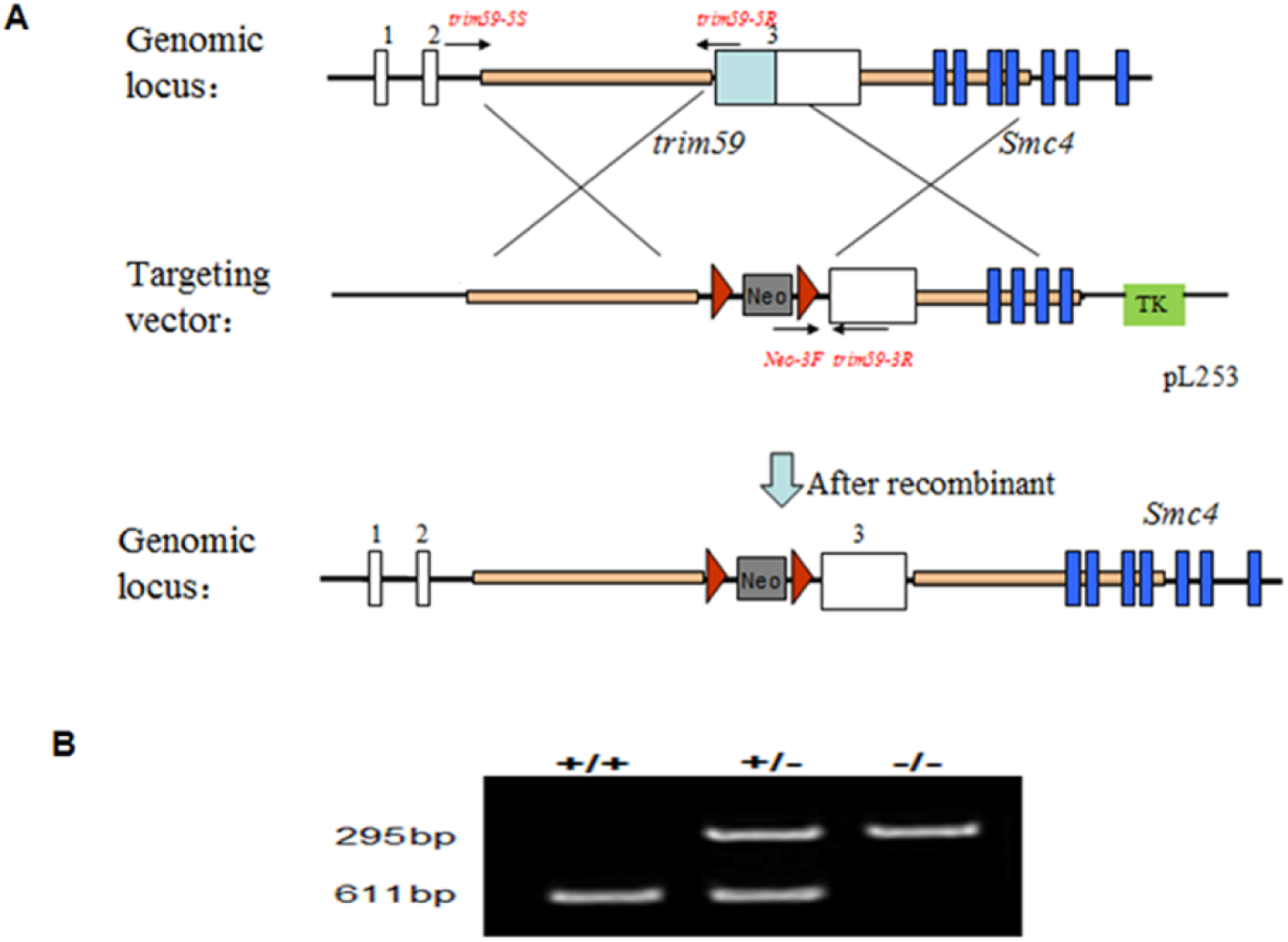
Generation and identification of *Trim59* knockout mice. (A) Strategy for generating *Trim59* KO mice. *Trim59* KO mice were generated according to the described protocol in materials and methods. (B) PCR of mouse tail genome DNA. Total tail genome DNA was extracted using a DNA kit and PCR was performed. +/+, wild-type mouse; +/-, heterozygote mouse; -/-, *Trim59* KO mouse.

**Fig. S3.**
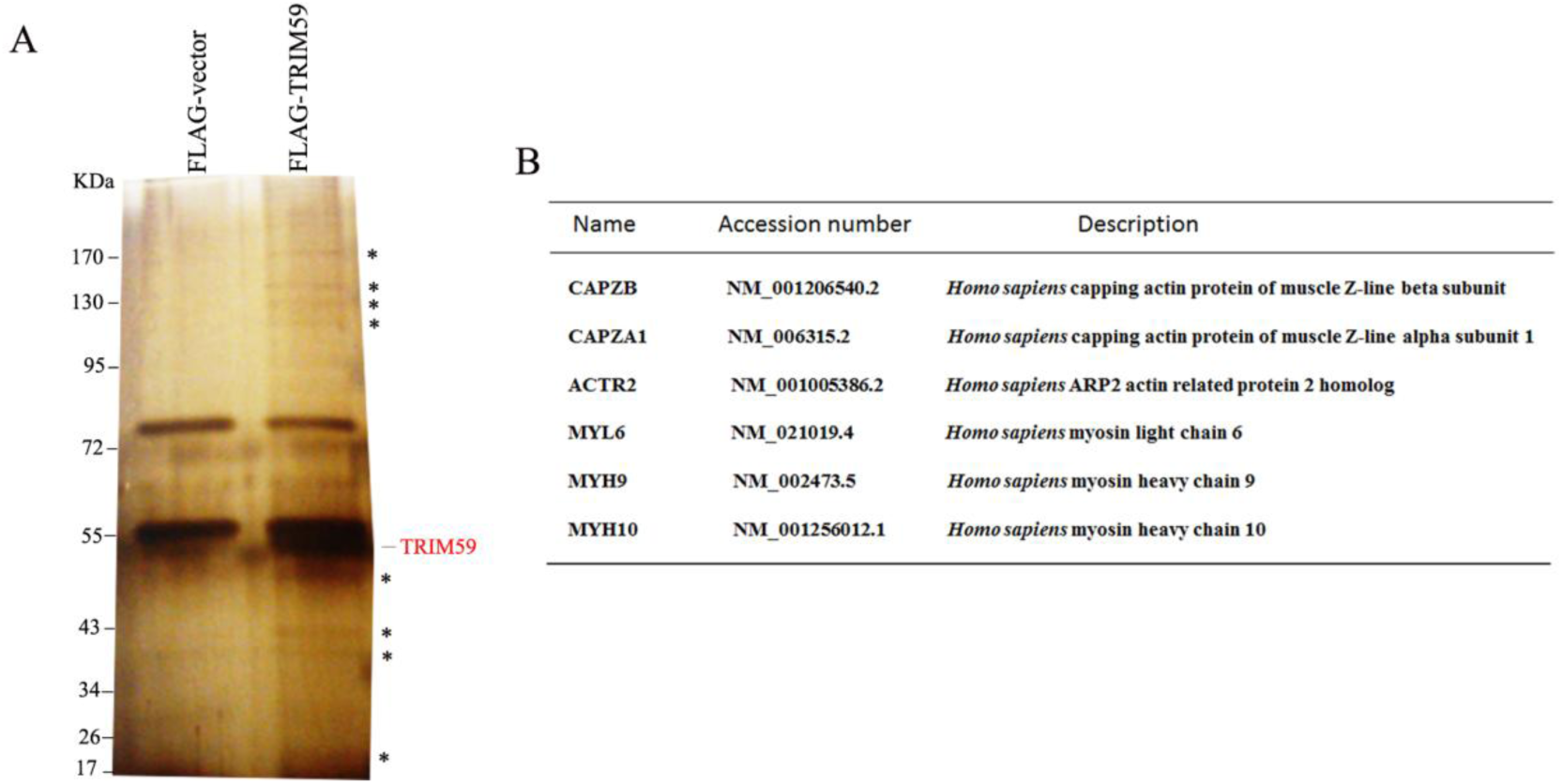
Interaction of Trim59 and actin-associated proteins. (A) Immunoprecipitation analyses. HEK293T cells were transfected with FLAG-vector or FLAG-Trim59. After immunoprecipitation by anti-FLAG, proteins were separated by SDS-PAGE, and then stained using silver staining kit. Asterisks indicate proteins which only appeared in Trim59 overexpressing group. (B) Proteins which are relative to the functions of cytoskeleton which only appeared in FLAG-Trim59 overexpressing group. The gel lanes containing the immunopurified samples were excised for Liquid Chromatography-tandem Mass (LC-MS/MS) analysis.

**Fig. S4.**
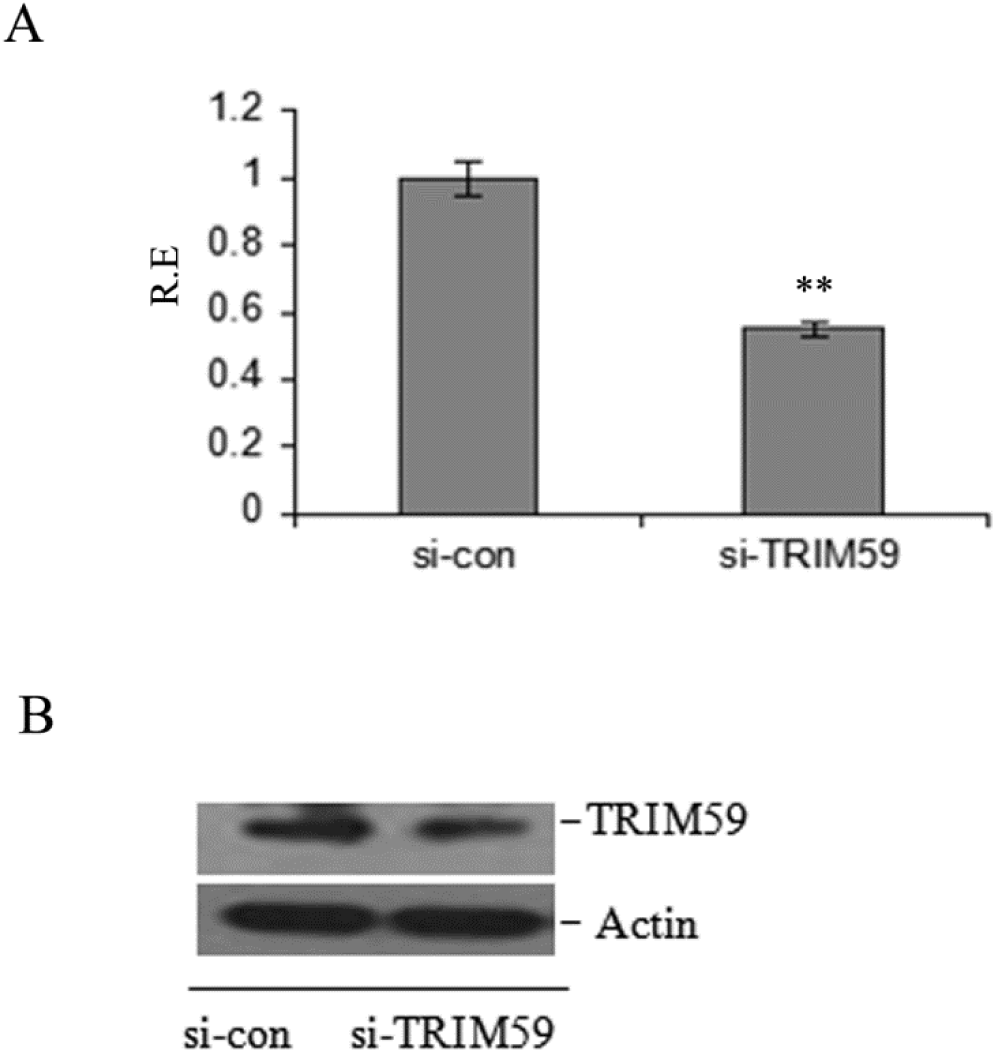
Expression of Trim59 in siRNA silencing mouse embryonic stem cells. (A) qRT-PCR of Trim59 in the F1 ESCs with (si-Trim59) or without (si-Con) mouse Trim59 siRNA treatment. ^∗^*p*<0.05 and ^∗∗^*p*<0.01(t-test). (B) Immunoblot of Trim59 in the F1 ESCs with (si-Trim59) or without (si-Con) mouse Trim59 siRNA treatment. Actin, a loading control.

**Fig. S5.**
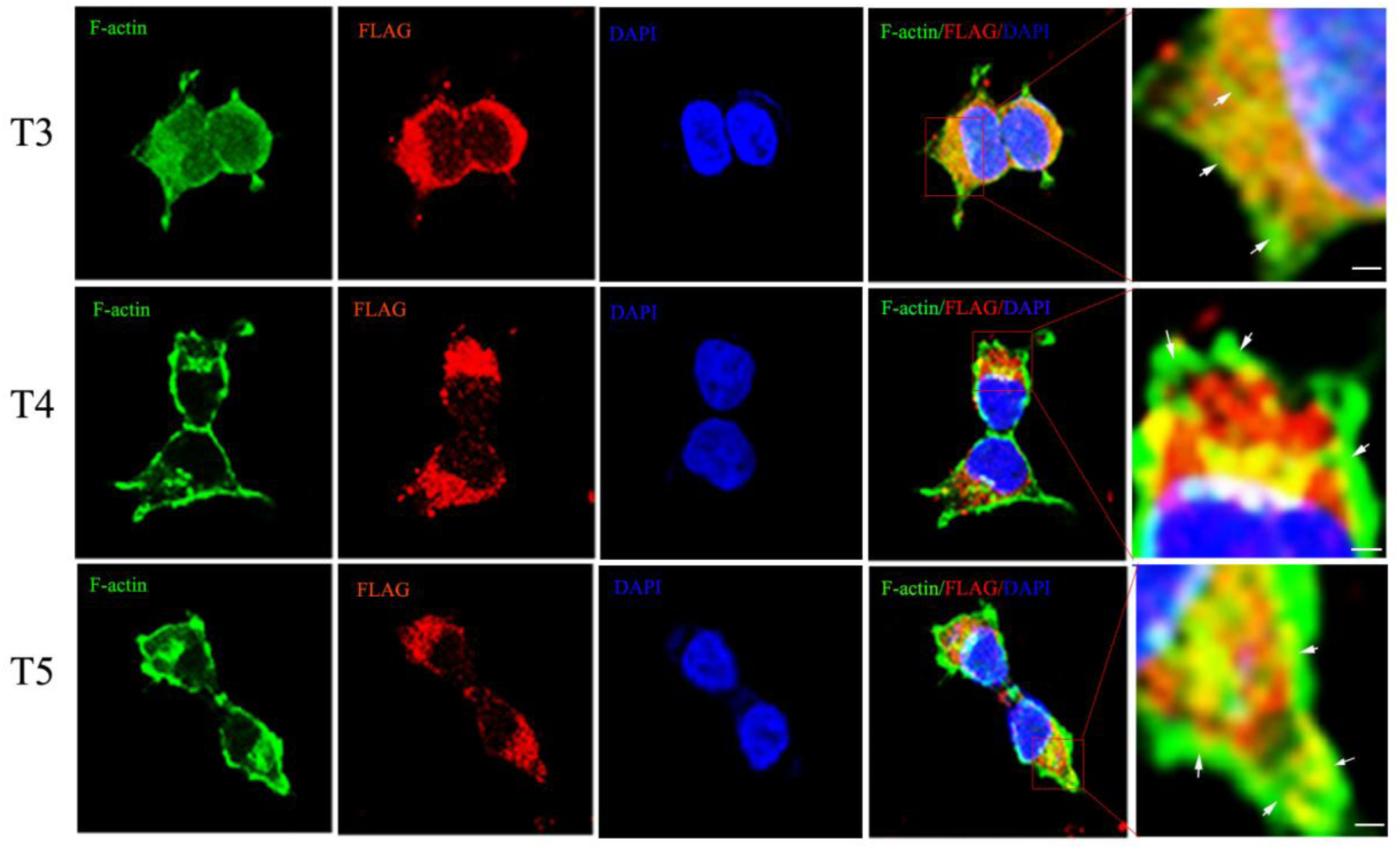
Trim59 RING-finger domain is required for F-actin assembly. HEK293T cells were transfected with expression vectors encoding FLAG-tagged Trim59 domain mutants for 24hrs. The expression of FLAG-tag Trim59 domain deleted mutants (T3, T4, T5) and the F-actin assembly were analyzed by immunofluorescence. F-actin (green), Trim59 mutants (red) and chromatin (blue). Images were acquired using a Bio-Rad Radiance 2100 confocal microscope with a Zeiss 63× oil immersion objective; scale bar, 10 μm; zoom: ×4.

**Fig. S6.**
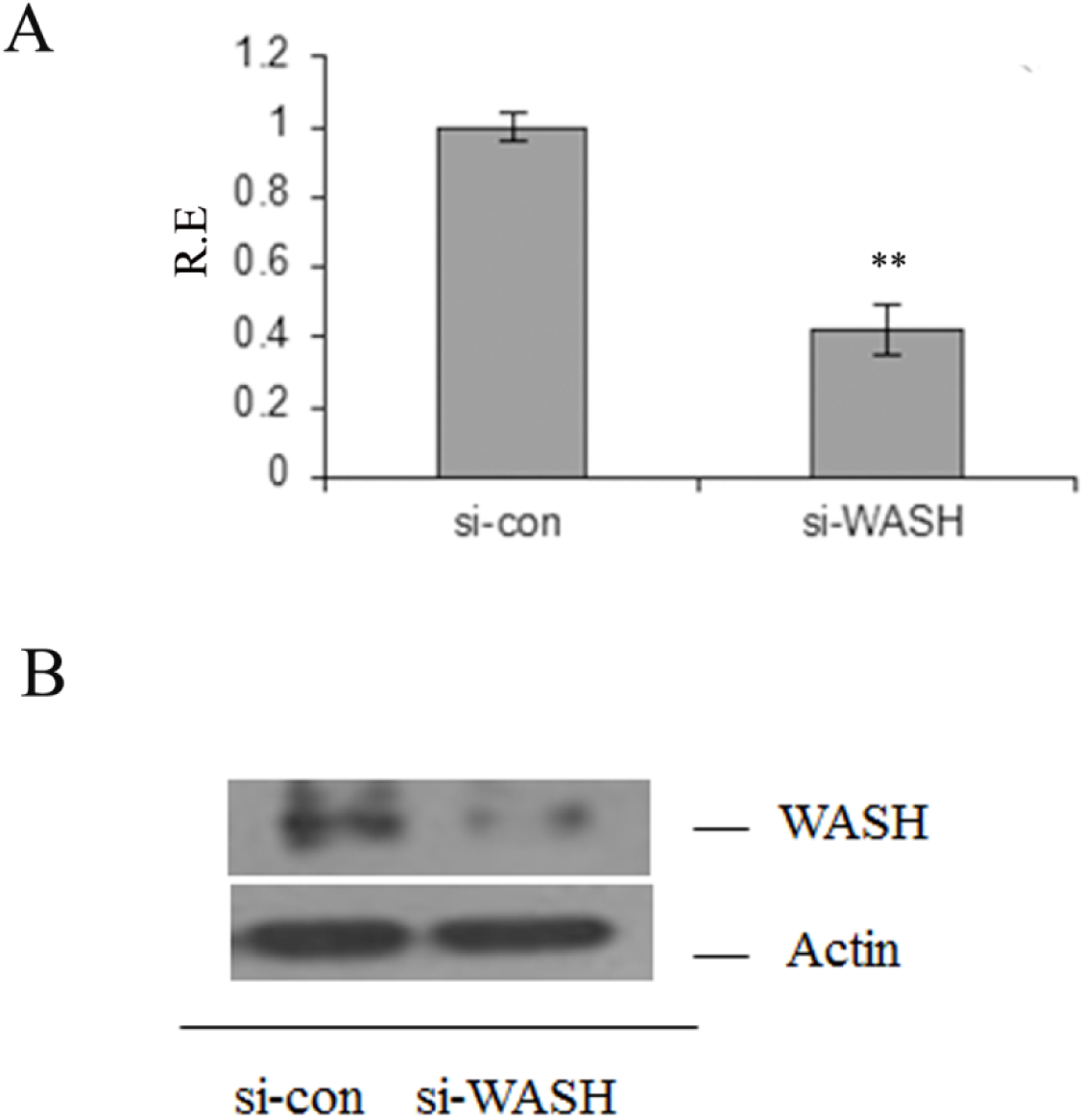
Expression of WASH after siRNA treatment. (A) qRT-PCR of WASH in F1 ESCs with (si-WASH) or without (si-Con) mouse WASH siRNA.^∗^*p*<0.05 and ^∗∗^*p*<0.01(t-test). (B) Immunoblot of WASH in F1 ESCs with (si-WASH) or without (si-Con) mouse WASH siRNA. Actin, a loading control.

**Fig. S7.**
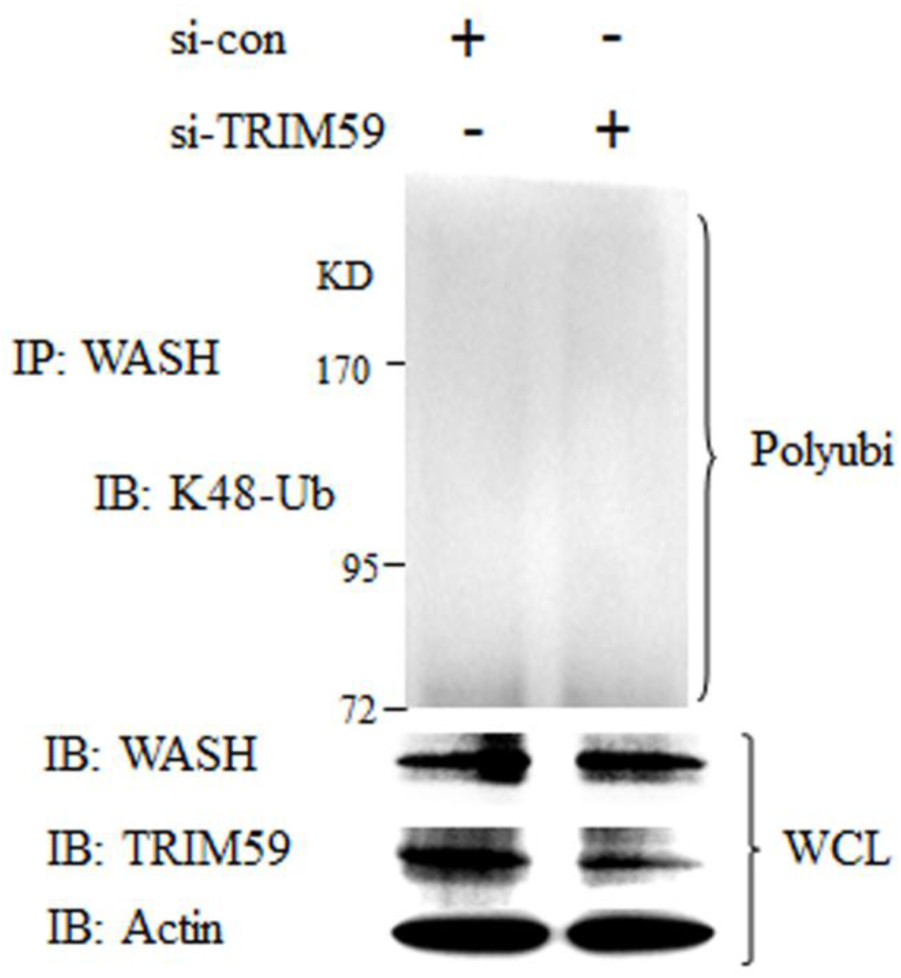
Immunoblot of K48-linked ubiquitination of endogenous WASH. Immunoblot analysis of K48-linked ubiquitination (K48-Ub) of endogenouse WASH which was immunoprecipitated in F1 ESCs after treatent with mouse Trim59 siRNA (top blot). Trim59, WASH and actin (below blots) were detected in the same cells without immunoprecipitation. Polyubi., polyubiquitination; IP, immunoprecipitation; IB, immunoblot assay; Actin, a loading control; WCL, whole cell lysates.

## Supplementary Table S3

**Table S3A.**
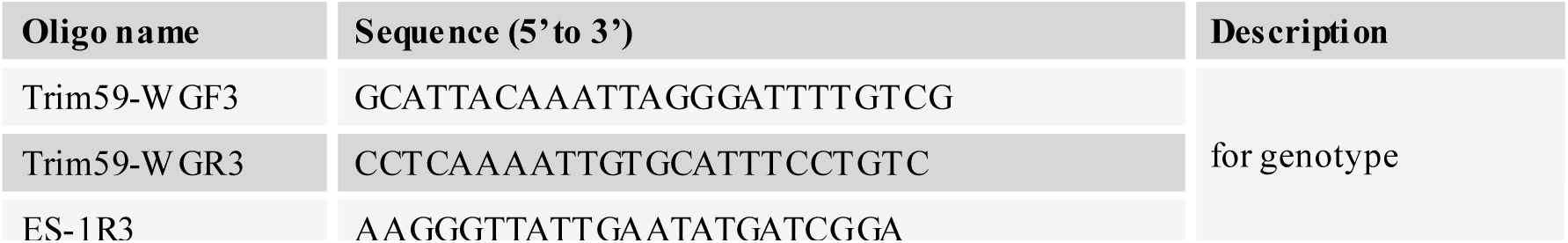
Oligoes used in the identification of mouse genotype.

**Table S3B.**
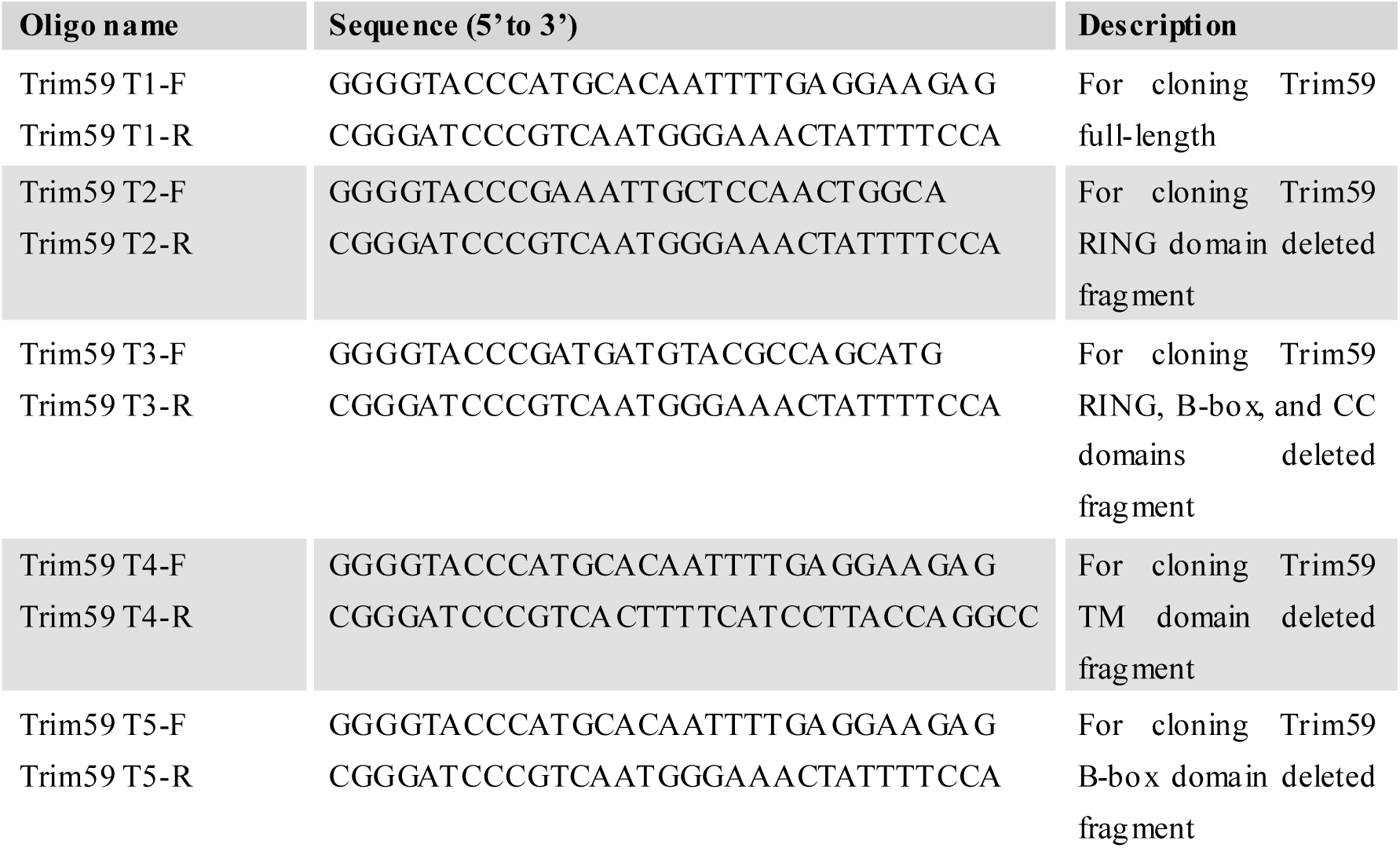
Primers used in cloning Trim59 full-length and mutants.

**Table S3C.**
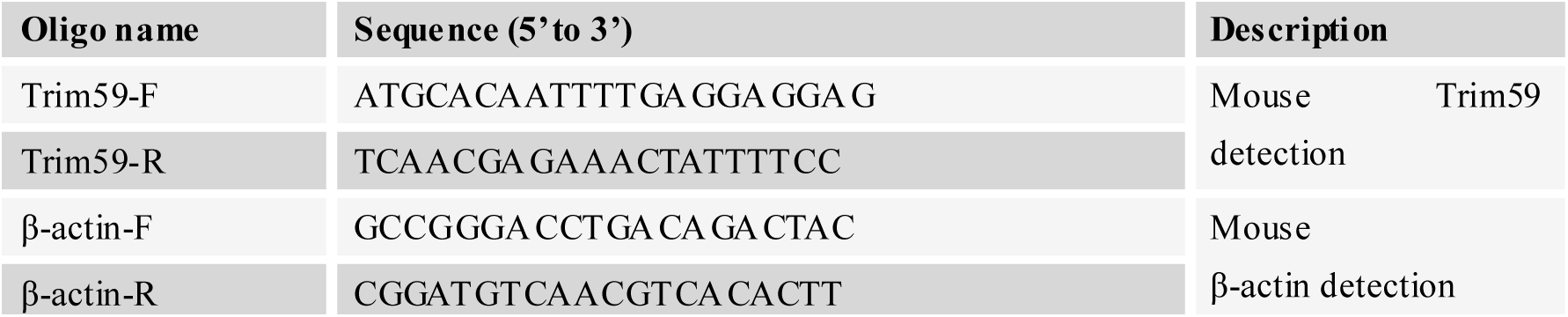
Oligoes used in RT-PCR.

**Table S3D.**
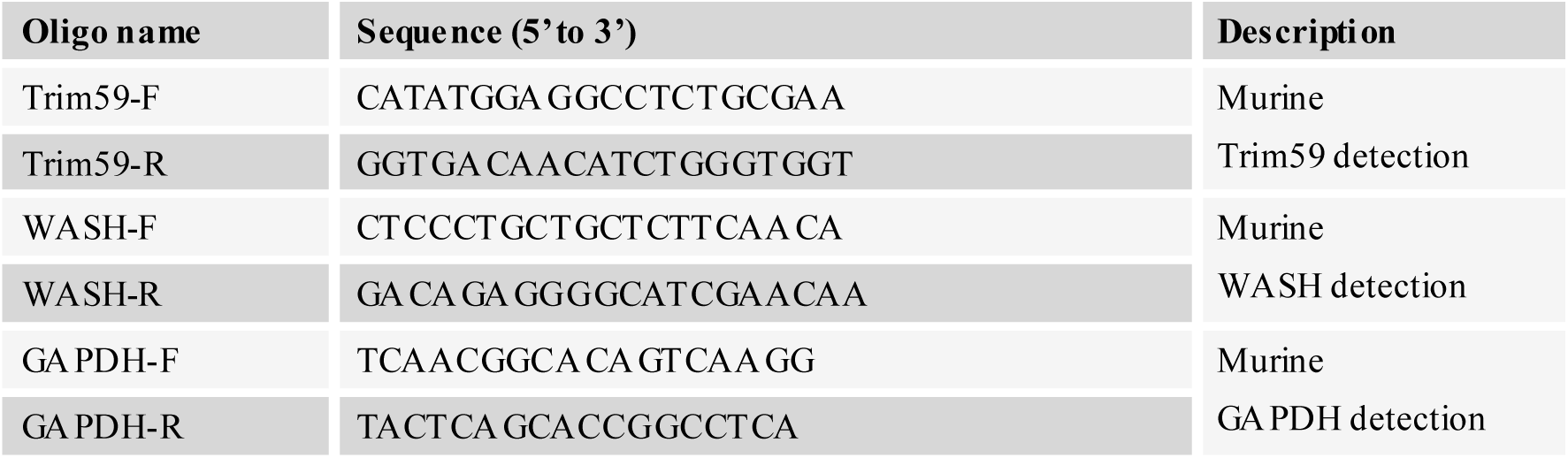
Oligoes used in qPCR.

**Table S3E.**
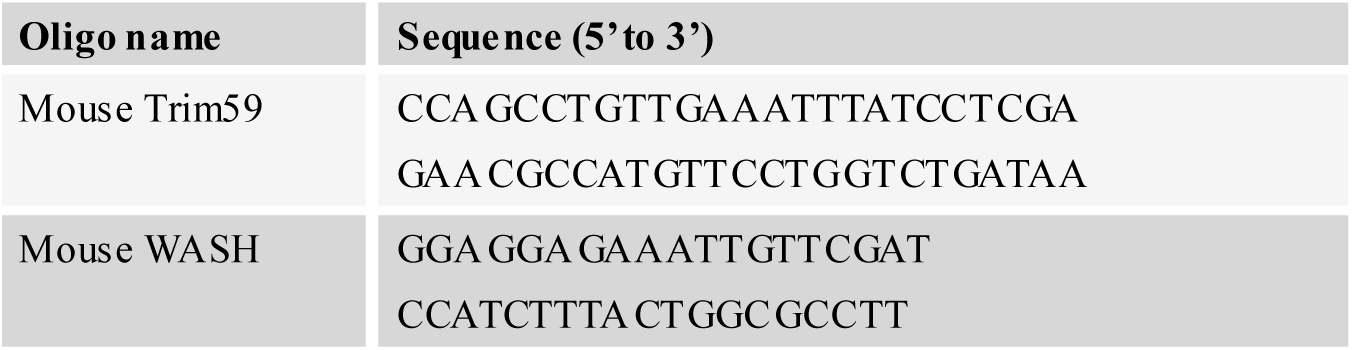
SiRNA sequences used in this study.

**Table S3F.**
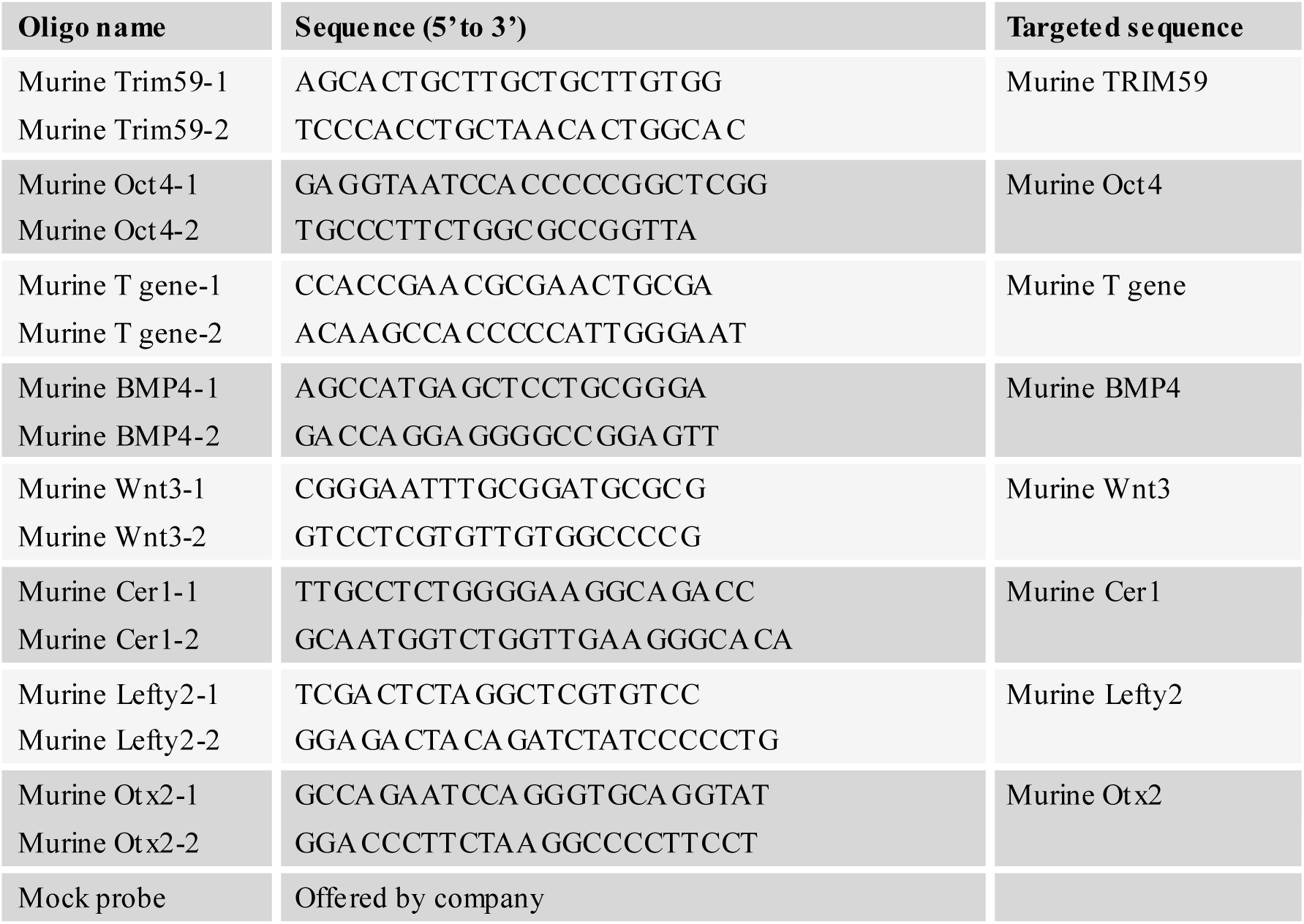
Probes used in *in situ* hybridization.

## Reference

Aierken, G., Seyiti, A., Alifu, M., and Kuerban, G. (2017). Knockdown of Tripartite-59 (TRIM59) Inhibits Cellular Proliferation and Migration in Human Cervical Cancer Cells. Oncology Research 25, 381-388.

Arias, A. M., and Hayward, P. (2006). Filtering transcriptional noise during development: concepts and mechanisms. Nature Reviews Genetics 7, 34-44.

Barrow, J. R., Howell, W. D., Rule, M., Hayashi, S., Thomas, K. R., Capecchi, M. R., and McMahon, A. R. (2007). Wnt3 signaling in the epiblast is required for proper orientation of the anteroposterior axis. Developmental Biology 312, 312-320.

Basilicata, M. F., Frank, M., Solter, D., Brabletz, T., and Stemmler, M. P. (2016). Inappropriate cadherin switching in the mouse epiblast compromises proper signaling between the epiblast and the extraembryonic ectoderm during gastrulation. Scientific Reports 6.

Bassham, S., and Postlethwait, J. (2000). Brachyury (T) expression in embryos of a larvacean urochordate, Oikopleura dioica, and the ancestral role of T. Dev Biol 220, 322-332.

Blitz, I. L., Cho, K. W. Y., and Chang, C. B. (2003). Twisted gastrulation loss-of-function analyses support its role as a BMP inhibitor during early Xenopus ernbryogenesis. Development 130, 4975-4988.

Boiani, M., Eckardt, S., Scholer, H. R., and McLaughlin, K. J. (2002). Oct4 distribution and level in mouse clones: consequences for pluripotency. Genes Dev 16, 1209-1219.

Boumela, I., Assou, S., Aouacheria, A., Haouzi, D., Dechaud, H., De Vos, J., Handyside, A., and Hamamah, S. (2011). Involvement of BCL2 family members in the regulation of human oocyte and early embryo survival and death: gene expression and beyond. Reproduction 141, 549-561.

Campellone, K. G., and Welch, M. D. (2010). A nucleator arms race: cellular control of actin assembly. Nature reviews Molecular cell biology 11, 237-251.

Caronna, E. A., Patterson, E. S., Hummert, P. M., and Kroll, K. L. (2013). Geminin Restrains Mesendodermal Fate Acquisition of Embryonic Stem Cells and is Associated with Antagonism of Wnt Signaling and Enhanced Polycomb-Mediated Repression. Stem cells 31, 1477-1487.

Chazaud, C., and Rossant, J. (2006). Disruption of early proximodistal patterning and AVE formation in Apc mutants. Development 133, 3379-3387.

Chen, W., Zhao, K., Miao, C. K., Xu, A. M., Zhang, J. Z., Zhu, J. D., Su, S. F., and Wang, Z. J. (2017). Silencing Trim59 inhibits invasion/migration and epithelial-to-mesenchymal transition via TGF-beta/Smad2/3 signaling pathway in bladder cancer cells. Oncotargets and Therapy 10, 1503-1512.

Cuevas, E., Rybak-Wolf, A., Rohde, A. M., Nguyen, D. T., and Wulczyn, F. G. (2015). Lin41/Trim71 is essential for mouse development and specifically expressed in postnatal ependymal cells of the brain. Front Cell Dev Biol 3, 20.

De Mot, L., Gonze, D., Bessonnard, S., Chazaud, C., Goldbeter, A., and Dupont, G. (2016). Cell Fate Specification Based on Tristability in the Inner Cell Mass of Mouse Blastocysts. Biophys J 110, 710-722.

Deng, L., Wang, C., Spencer, E., Yang, L., Braun, A., You, J., Slaughter, C., Pickart, C., and Chen, Z. J. (2000). Activation of the IkappaB kinase complex by TRAF6 requires a dimeric ubiquitin-conjugating enzyme complex and a unique polyubiquitin chain. Cell 103, 351-361.

Derivery, E., Sousa, C., Gautier, J. J., Lombard, B., Loew, D., and Gautreau, A. (2009). The Arp2/3 activator WASH controls the fission of endosomes through a large multiprotein complex. Developmental cell 17, 712-723.

Esposito, D., Koliopoulos, M. G., and Rittinger, K. (2017). Structural determinants of TRIM protein function. Biochem Soc Trans 45, 183-191.

Gaivao, M. M. F., Rambags, B. P. B., and Stout, T. A. E. (2014). Gastrulation and the establishment of the three germ layers in the early horse conceptus. Theriogenology 82, 354-365.

Goncalves, L., Filipe, M., Marques, S., Salgueiro, A. M., Becker, J. D., and Belo, J. A. (2011a). Identification and functional analysis of novel genes expressed in the Anterior Visceral Endoderm. Int J Dev Biol 55, 281-295.

Goncalves, L., Filipe, M., Marques, S., Salgueiro, A. M., Becker, J. D., and Belo, J. A. (2011b). Identification and functional analysis of novel genes expressed in the Anterior Visceral Endoderm. International Journal of Developmental Biology 55, 281-295.

Hao, Y. H., Doyle, J. M., Ramanathan, S., Gomez, T. S., Jia, D., Xu, M., Chen, Z. J., Billadeau, D. D., Rosen, M. K., and Potts, P. R. (2013). Regulation of WASH-dependent actin polymerization and protein trafficking by ubiquitination. Cell 152, 1051-1064.

Herrmann, B. G. (1991). Expression pattern of the Brachyury gene in whole-mount TWis/TWis mutant embryos. Development 113, 913-917.

Hoshino, H., Shioi, G., and Aizawa, S. (2015). AVE protein expression and visceral endoderm cell behavior during anterior-posterior axis formation in mouse embryos: Asymmetry in OTX2 and DKK1 expression. Dev Biol 402, 175-191.

Houston, D. W., and Cuykendall, T. N. (2009). Localized Xenopus Trim36 regulates cortical rotation and dorsal axis formation. Developmental Biology 331, 391-391.

James, L. C., Keeble, A. H., Khan, Z., Rhodes, D. A., and Trowsdale, J. (2007). Structural basis for PRYSPRY-mediated tripartite motif (TRIM) protein function. Proceedings of the National Academy of Sciences of the United States of America 104, 6200-6205.

Jia, D., Gomez, T. S., Metlagel, Z., Umetani, J., Otwinowski, Z., Rosen, M. K., and Billadeau, D. D. (2010). WASH and WAVE actin regulators of the Wiskott-Aldrich syndrome protein (WASP) family are controlled by analogous structurally related complexes. Proceedings of the National Academy of Sciences of the United States of America 107, 10442-10447.

Joubin, K., and Stern, C. D. (1999). Molecular interactions continuously define the organizer during the cell movements of gastrulation. Cell 98, 559-571.

Ke, H., Parron, V. I., Reece, J., Zhang, J. Y., Akiyama, S. K., and French, J. E. (2010). BCL2 inhibits cell adhesion, spreading, and motility by enhancing actin polymerization. Cell Res 20, 458-469.

Kelly, O. G., Pinson, K. I., and Skarnes, W. C. (2004). The Wnt co-receptors Lrp5 and Lrp6 are essential for gastrulation in mice. Development 131, 2803-2815.

Khatamianfar, V., Valiyeva, F., Rennie, P. S., Lu, W. Y., Yang, B. B., Bauman, G. S., Moussa, M., and Xuan, J. W. (2012). TRIM59, a novel multiple cancer biomarker for immunohistochemical detection of tumorigenesis. Bmj Open 2.

Kim, D. K., Cha, Y., Ahn, H. J., Kim, G., and Park, K. S. (2014). Lefty1 and Lefty2 Control the Balance Between Self-Renewal and Pluripotent Differentiation of Mouse Embryonic Stem Cells. Stem Cells and Development 23, 457-466.

Kim, J., and Kaartinen, V. (2008). Generation of mice with a conditional allele for Trim33. Genesis 46, 329-333.

Kitamura, K., Tanaka, H., and Nishimune, Y. (2005). The RING-finger protein haprin: domains and function in the acrosome reaction. Curr Protein Pept Sci 6, 567-574.

Kojima, Y., Tam, O. H., and Tam, P. P. L. (2014). Timing of developmental events in the early mouse embryo. Seminars in Cell & Developmental Biology 34, 65-75.

Kurek, D., Neagu, A., Tastemel, M., Tuysuz, N., Lehmann, J., van de Werken, H. J. G., Philipsen, S., van der Linden, R., Maas, A., van IJcken, W. F. J., et al. (2015). Endogenous WNT Signals Mediate BMP-Induced and Spontaneous Differentiation of Epiblast Stem Cells and Human Embryonic Stem Cells. Stem Cell Reports 4, 114-128.

Lanier, L. M., Gates, M. A., Witke, W., Menzies, A. S., Wehman, A. M., Macklis, J. D., Kwiatkowski, D., Soriano, P., and Gertler, F. B. (1999). Mena is required for neurulation and commissure formation. Neuron 22, 313-325.

Le Bin, G. C., Munoz-Descalzo, S., Kurowski, A., Leitch, H., Lou, X., Mansfield, W., Etienne-Dumeau, C., Grabole, N., Mulas, C., Niwa, H., et al. (2014). Oct4 is required for lineage priming in the developing inner cell mass of the mouse blastocyst. Development 141, 1001-1010.

Lenhart, K. F., Lin, S. Y., Titus, T. A., Postlethwait, J. H., and Burdine, R. D. (2011). Two additional midline barriers function with midline lefty1 expression to maintain asymmetric Nodal signaling during left-right axis specification in zebrafish. Development 138, 4405-4410.

Li, Y., Wu, H., Wu, W., Zhuo, W., Liu, W., Zhang, Y., Cheng, M., Chen, Y. G., Gao, N., Yu, H., et al. (2014). Structural insights into the TRIM family of ubiquitin E3 ligases. Cell Res 24, 762-765.

Liguori, G. L., Borges, A. C., D’Andrea, D., Liguoro, A., Goncalves, L., Salgueiro, A. M., Persico, M. G., and Belo, J. A. (2008). Cripto-independent Nodal signaling promotes positioning of the A-P axis in the early mouse embryo. Dev Biol 315, 280-289.

Liu, P., Wakamiya, M., Shea, M. J., Albrecht, U., Behringer, R. R., and Bradley, A. (1999). Requirement for Wnt3 in vertebrate axis formation. Nature genetics 22, 361-365.

Luo, D. K., Wang, Y. N., Huan, X. K., Huang, C., Yang, C., Fan, H., Xu, Z. K., and Yang, L. (2017). Identification of a synonymous variant in TRIM59 gene for gastric cancer risk in a Chinese population. Oncotarget 8, 11507-11516.

Meroni, G., and Diez-Roux, G. (2005). TRIM/RBCC, a novel class of ‘single protein RING finger’ E3 ubiquitin ligases. Bioessays 27, 1147-1157.

Montazeri, M., Sanchez-Lopez, J. A., Caballero, I., Lay, N. M., Elliott, S., Lopez-Martin, S., Yanez-Mo, M., and Fazeli, A. (2015). Activation of Toll-like receptor 3 reduces actin polymerization and adhesion molecule expression in endometrial cells, a potential mechanism for viral-induced implantation failure. Human Reproduction 30, 893-905.

Nichols, J., Zevnik, B., Anastassiadis, K., Niwa, H., Klewe-Nebenius, D., Chambers, I., Scholer, H., and Smith, A. (1998). Formation of pluripotent stem cells in the mammalian embryo depends on the POU transcription factor Oct4. Cell 95, 379-391.

Niedenberger, B. A., Chappell, V. A., Otey, C. A., and Geyer, C. B. (2014). Actin dynamics regulate subcellular localization of the F-actin-binding protein PALLD in mouse Sertoli cells. Reproduction 148, 333-341.

Niwa, H., Miyazaki, J., and Smith, A. G. (2000). Quantitative expression of Oct-3/4 defines differentiation, dedifferentiation or self-renewal of ES cells. Nature genetics 24, 372-376.

Padgett, R. W., Wozney, J. M., and Gelbart, W. M. (1993). Human BMP sequences can confer normal dorsal-ventral patterning in the Drosophila embryo. Proceedings of the National Academy of Sciences of the United States of America 90, 2905-2909.

Parfitt, D. E., and Shen, M. M. (2014). From blastocyst to gastrula: gene regulatory networks of embryonic stem cells and early mouse embryogenesis. Philos Trans R Soc Lond B Biol Sci 369.

Pesce, M., and Scholer, H. R. (2001). Oct-4: gatekeeper in the beginnings of mammalian development. Stem cells 19, 271-278.

Ray, R. P., Arora, K., Nusslein-Volhard, C., and Gelbart, W. M. (1991). The control of cell fate along the dorsal-ventral axis of the Drosophila embryo. Development 113, 35-54.

Sasaki, H. (2017). Roles and regulations of Hippo signaling during preimplantation mouse development. Development, growth & differentiation 59, 12-20.

Simeone, A., and Acampora, D. (2001). The role of Otx2 in organizing the anterior patterning in mouse. International Journal of Developmental Biology 45, 337-345.

Soares, M. L., Torres-Padilla, M. E., and Zernicka-Goetz, M. (2008). Bone morphogenetic protein 4 signaling regulates development of the anterior visceral endoderm in the mouse embryo. Development, growth & differentiation 50, 615-621.

Spiller, C. M., Feng, C. W., Jackson, A., Gillis, A. J. M., Rolland, A. D., Looijenga, L. H. J., Koopman, P., and Bowles, J. (2012). Endogenous Nodal signaling regulates germ cell potency during mammalian testis development. Development 139, 4123-4132.

Sun, L., and Chen, Z. J. (2004). The novel functions of ubiquitination in signaling. Current opinion in cell biology 16, 119-126.

Takenaga, M., Fukumoto, M., and Hori, Y. (2007). Regulated Nodal signaling promotes differentiation of the definitive endoderm and mesoderm from ES cells. J Cell Sci 120, 2078-2090.

Tam, P. P. L., Goldman, D., Camus, A., and Schoenwolf, G. C. (2000). Early events of somitogenesis in higher vertebrates: Allocation of precursor cells during gastrulation and the organization of a meristic pattern in the paraxial mesoderm. Current Topics in Developmental Biology, Vol 47 47, 1-32.

Tan, K., An, L., Wang, S. M., Wang, X. D., Zhang, Z. N., Miao, K., Sui, L. L., He, S. Z., Nie, J. Z., Wu, Z. H., and Tian, J. H. (2015a). Actin Disorganization Plays a Vital Role in Impaired Embryonic Development of In Vitro-Produced Mouse Preimplantation Embryos. Plos One 10, e0130382.

Tan, K., An, L., Wang, S. M., Wang, X. D., Zhang, Z. N., Miao, K., Sui, L. L., He, S. Z., Nie, J. Z., Wu, Z. H., and Tian, J. H. (2015b). Actin Disorganization Plays a Vital Role in Impaired Embryonic Development of In Vitro-Produced Mouse Preimplantation Embryos. Plos One 10.

Thomson, M., Liu, S. J., Zou, L. N., Smith, Z., Meissner, A., and Ramanathan, S. (2011). Pluripotency Factors in Embryonic Stem Cells Regulate Differentiation into Germ Layers. Cell 145, 875-889.

Wang, F., Zhang, L., Zhang, G. L., Wang, Z. B., Cui, X. S., Kim, N. H., and Sun, S. C. (2014). WASH complex regulates Arp2/3 complex for actin-based polar body extrusion in mouse oocytes. Sci Rep 4, 5596.

Wilcockson, S. G., Sutcliffe, C., and Ashe, H. L. (2017). Control of signaling molecule range during developmental patterning. Cellular and Molecular Life Sciences 74, 1937-1956.

Xia, P., Wang, S., Du, Y., Zhao, Z., Shi, L., Sun, L., Huang, G., Ye, B., Li, C., Dai, Z., et al. (2013). WASH inhibits autophagy through suppression of Beclin 1 ubiquitination. EMBO J 32, 2685-2696.

Yan, J., Li, Q., Mao, A. P., Hu, M. M., and Shu, H. B. (2014). TRIM4 modulates type I interferon induction and cellular antiviral response by targeting RIG-I for K63-linked ubiquitination. J Mol Cell Biol 6, 154-163.

Yoon, Y., Huang, T. T., Tortelote, G. G., Wakamiya, M., Hadjantonakis, A. K., Behringer, R. R., and Rivera-Perez, J. A. (2015). Extra-embryonic Wnt3 regulates the establishment of the primitive streak in mice. Developmental Biology 403, 80-88.

Zhang, Y., and Yang, W. B. (2017). Down-regulation of tripartite motif protein 59 inhibits proliferation, migration and invasion in breast cancer cells. Biomedicine & Pharmacotherapy 89, 462-467.

Zhang, Z., Zeng, B., Zhang, Z., Jiao, G., Li, H., Jing, Z., Ouyang, J., Yuan, X., Chai, L., Che, Y., et al. (2009). Suppressor of cytokine signaling 3 promotes bone marrow cells to differentiate into CD8+ T lymphocytes in lung tissue via up-regulating Notch1 expression. Cancer Res 69, 1578-1586.

Zhao, X. F., Liu, Q. H., Du, B. Q., Li, P., Cui, Q., Han, X., Du, B. R., Yan, D. M., and Zhu, X. (2012). A novel accessory molecule Trim59 involved in cytotoxicity of BCG-activated macrophages. Molecules and Cells 34, 263-270.

Zhou, Z., Ji, Z., Wang, Y., Li, J., Cao, H., Zhu, H. H., and Gao, W. Q. (2014). TRIM59 is up-regulated in gastric tumors, promoting ubiquitination and degradation of p53. Gastroenterology 147, 1043-1054.

